# Kv2.1 mediates spatial and functional coupling of L-type calcium channels and ryanodine receptors in neurons

**DOI:** 10.1101/702514

**Authors:** Nicholas C. Vierra, Michael Kirmiz, Deborah van der List, L. Fernando Santana, James S. Trimmer

**Affiliations:** Department of Neurobiology, Physiology, and Behavior, University of California, Davis, CA 95616; Department of Physiology and Membrane Biology, University of California, Davis, School of Medicine, Davis, CA 95616

## Abstract

The voltage-gated K^+^ channel Kv2.1 serves a major structural role in the soma and proximal dendrites of brain neurons, tethering the plasma membrane (PM) to the endoplasmic reticulum (ER). Although Kv2.1 clustering at neuronal ER-PM junctions (EPJs) is tightly regulated and conserved across species, its function at these sites is unclear. By identifying and evaluating proteins in close spatial proximity to Kv2.1-containing EPJs, we discovered that a significant role of Kv2.1 at EPJs is to promote the clustering and functional coupling of PM L-type Ca^2+^ channels (LTCCs) to ryanodine receptor (RyR) ER Ca^2+^ release channels. Kv2.1 clustering also unexpectedly enhanced LTCC opening at polarized membrane potentials. This enabled Kv2.1-LTCC-RyR triads to generate localized Ca^2+^ release events (i.e., Ca^2+^ sparks) independently of action potentials. Together, these findings uncover a novel mode of LTCC regulation and establish a unique mechanism whereby Kv2.1-associated EPJs provide a molecular platform for localized somatodendritic Ca^2+^ signals.

## Introduction

The members of the Kv2 family of voltage-gated K^+^ (Kv) channels, Kv2.1 and Kv2.2, are among the most abundant and widely expressed K^+^ channels in mammalian brain neurons (Trimmer, 2015). Kv2 channels are present in high density clusters [approximately 3.7-fold greater channel density within clusters than in the adjacent membrane (Fox et al., 2013)] localized to neuronal somata, proximal dendrites, and axon initial segments (Trimmer, 1991; Du et al., 1998; Bishop et al., 2015; Kirmiz et al., 2018a). In hippocampal and cortical neurons, Kv2 channels conduct most of the delayed rectifier K^+^ current (Murakoshi and Trimmer, 1999; Du et al., 2000; Guan et al., 2007). Detailed studies have revealed the significant influence of neuronal Kv2.1-mediated currents on action potential duration and repetitive firing (Du et al., 2000; Liu and Bean, 2014; Kimm et al., 2015). In addition to its important role in modulating intrinsic electrical activity, Kv2.1 serves a non-canonical structural (*i.e.*, non-conducting) function in tethering the plasma membrane (PM) to the endoplasmic reticulum (ER) to form ER-PM junctions (EPJs) (Fox et al., 2015; Johnson et al., 2018; Kirmiz et al., 2018a; Kirmiz et al., 2018b). Although Kv2.1 clustering at EPJs is tightly regulated and independent of K^+^ conductance (Kirmiz et al., 2018a), the physiological impact of concentrating this Kv channel at an EPJ is not known.

In brain neurons, EPJs occupy approximately 10% of the PM surface area, predominantly within the soma and proximal dendrites (Wu et al., 2017). By electron microscopy, the ER at many neuronal EPJs appears as a micron-diameter, flattened vesicle less than 10 nm from the PM, designated a subsurface cistern (Rosenbluth, 1962; Tao-Cheng, 2018). While the specific functions of neuronal subsurface cisterns remain unclear, in most eukaryotic cells, EPJs represent domains specialized for maintenance of Ca^2+^, lipid, and metabolic homeostasis (Gallo et al., 2016; Chang et al., 2017).

L-type voltage-gated Ca^2+^ channels (LTCCs) are prominently expressed in neurons throughout the brain (Catterall, 2011; Zamponi et al., 2015). Their important role in brain is underscored by studies showing genetic variation in the *CACNA1C* gene encoding Cav1.2, the major voltage-sensing and pore forming α1 subunit expressed in brain, is associated with neurodevelopmental, psychiatric and neurological disorders (Splawski et al., 2004; Ferreira et al., 2008; Bozarth et al., 2018). Given their diverse and crucial roles in neuronal function, LTCCs are subjected to multimodal regulation to ensure their activity is coupled to overall cellular state especially as related to intracellular [Ca^2+^] (Lipscombe et al., 2013; Hofmann et al., 2014; Neely and Hidalgo, 2014). In both neurons and non-neuronal cells, Cav1.2-containing LTCCs are clustered at specific sites on the PM where they participate in supramolecular protein complexes that couple LTCC-mediated Ca^2+^ entry to specific Ca^2+^ signaling pathways (Dai et al., 2009; Rougier and Abriel, 2016). In neurons, LTCCs in dendritic spines participate in a complex whose output contributes to short- and long-term synaptic plasticity (Da Silva et al., 2013; Simms and Zamponi, 2014; Stanika et al., 2015; Wiera et al., 2017). Neocortical and hippocampal pyramidal neurons and dentate granule cells also have substantial LTCC populations in the soma and proximal dendrites (Westenbroek et al., 1990; Hell et al., 1993; Tippens et al., 2008; Berrout and Isokawa, 2009; Marshall et al., 2011; Kramer et al., 2012) representing the “aspiny” regions (Spruston and McBain, 2007) of these neurons. Many current models of Ca^2+^-dependent activation of transcription factors posit that somatic LTCCs uniquely contribute to transcription factor activation by mediating Ca^2+^ influx within specialized and compartmentalized signaling complexes (Wheeler et al., 2008; Ma et al., 2012; Matamales, 2012; Wheeler et al., 2012; Ma et al., 2014; Cohen et al., 2015; Yap and Greenberg, 2018; Wild et al., 2019). Yet, relatively little research has focused on the molecular mechanisms underlying the spatial and functional compartmentalization of the prominent somatic population of LTCCs compared to those on dendrites and at synapses.

Neuronal somata lack PM compartments analogous to dendritic spines, and fundamental questions remain as to how discrete Ca^2+^ signaling events can occur in the absence of such compartmentalization. In many non-neuronal cells, LTCCs are clustered at EPJs that represent specialized microdomains for LTCC-dependent and -independent Ca^2+^ signaling (Helle et al., 2013; Lam and Galione, 2013; Burgoyne et al., 2015; Henne et al., 2015; Gallo et al., 2016; Chung et al., 2017; Dickson, 2017). For example, Cav1.2-mediated Ca^2+^ entry is spatially and functionally coupled to ER ryanodine receptor (RyR) Ca^2+^ release channels at EPJs constituting the cardiomyocyte junctional dyad (Shuja and Colecraft, 2018). Localized Ca^2+^ release events (spreading <2 µm from the point of origin) called Ca^2+^ sparks arise from clusters of RyRs at these EPJs and are triggered *via* local Ca^2+^-induced Ca^2+^ release (CICR), a feed-forward phenomenon in which cytosolic Ca^2+^ binding to RyRs triggers their opening (Cheng et al., 1993; Cheng and Lederer, 2008). As indicated above, EPJs are abundant on neuronal somata (Wu et al., 2017), and neuronal somata have prominent LTCC- and RyR-mediated CICR (Friel and Tsien, 1992; Isokawa and Alger, 2006; Berrout and Isokawa, 2009). In addition, localized RyR-mediated Ca^2+^ release events occur in the somata and proximal dendrites of cultured and acute slice preparations of hippocampal pyramidal neurons (Koizumi et al., 1999; Berrout and Isokawa, 2009; Manita and Ross, 2009; Miyazaki et al., 2012), but a specific molecular structure underlying these events has not been described.

Given the well-characterized spatial and functional coupling of LTCCs and RyRs at EPJs in myocytes and previous observations of somatodendritic clustering of the LTCC Cav1.2 in hippocampal neurons (Westenbroek et al., 1990; Hell et al., 1993), our finding that Kv2.1 clusters are often juxtaposed to RyRs previously led us to hypothesize that Kv2.1 channels cluster with LTCCs to form Ca^2+^ “micro-signaling domains” (Antonucci et al., 2001; Misonou et al., 2005b). More recently, heterologously expressed Kv2.1 and Cav1.2 were found to colocalize in dissociated cultured hippocampal neurons (CHNs) (Fox et al., 2015). However, the spatial association of Kv2.1 with endogenous LTCCs and RyRs in brain neurons has not been determined. Here, we examined the subcellular distribution of Kv2.1, LTCCs, and RyRs in hippocampal neurons and used an unbiased proteomic analysis of brain tissue to identify LTCCs and RyRs as proteins in close spatial proximity to clustered Kv2.1. Using heterologous cells and CHNs, we investigated the impact of Kv2.1 clustering on the spatial coupling and functional properties of LTCCs and RyRs. We also defined how the localization and function of LTCCs and RyRs are affected by the loss of Kv2.1 in mouse CHNs lacking Kv2.1. Together, our findings establish a functional interaction between Kv2.1, LTCCs, and RyRs, reveal a significant influence of Kv2.1 in shaping neuronal LTCC activity, and support a critical role for Kv2.1 in the generation of somatodendritic Ca^2+^ signals.

## Results

### Kv2.1 channels spatially associate with LTCCs and RyRs in brain neurons

In mature CHNs, endogenous Cav1.2 channels are distributed to PM-localized clusters on the soma and proximal dendrites, distinct from their punctate localization in the more distal postsynaptic compartments that also contain the scaffolding protein PSD-95 (Di Biase et al., 2008) (Fig. 1A). To establish whether Kv2.1 channels spatially associate with Cav1.2, we examined rat CHNs immunolabeled for Kv2.1, Cav1.2, and RyRs. In the majority of CHNs expressing detectable levels of these proteins, presumed to be pyramidal neurons based on their morphological characteristics (Benson et al., 1994; Antonucci et al., 2001; Obermair et al., 2003), we observed overlapping clusters of Kv2.1 and RyRs that were spatially associated with smaller Cav1.2 clusters (Fig. 1B). We also observed more prominent spatial overlap of Cav1.2 and Kv2.1 immunolabeling in a subset of CHNs (Fig. 1C). Super-resolution structured illumination (SIM) imaging revealed that Kv2.1 clusters often encompassed smaller clusters of Cav1.2 as well Cav1.3 (Fig. 1D-E). We found that the spatial distributions of Kv2.1 and Cav1.2 puncta significantly correlated (p<0.001 versus the null hypothesis that the distributions of Kv2.1 and Cav1.2 puncta are independent) and could not be recapitulated in images in which their relative positions had been iteratively randomized in silico (Helmuth et al., 2010; Shivanandan et al., 2013). We also observed similar expression patterns of endogenous Cav1.3 and RyRs in CHNs, with Cav1.3 clusters spatially associated with RyR clusters (Fig. 1F).

**Figure 1.**
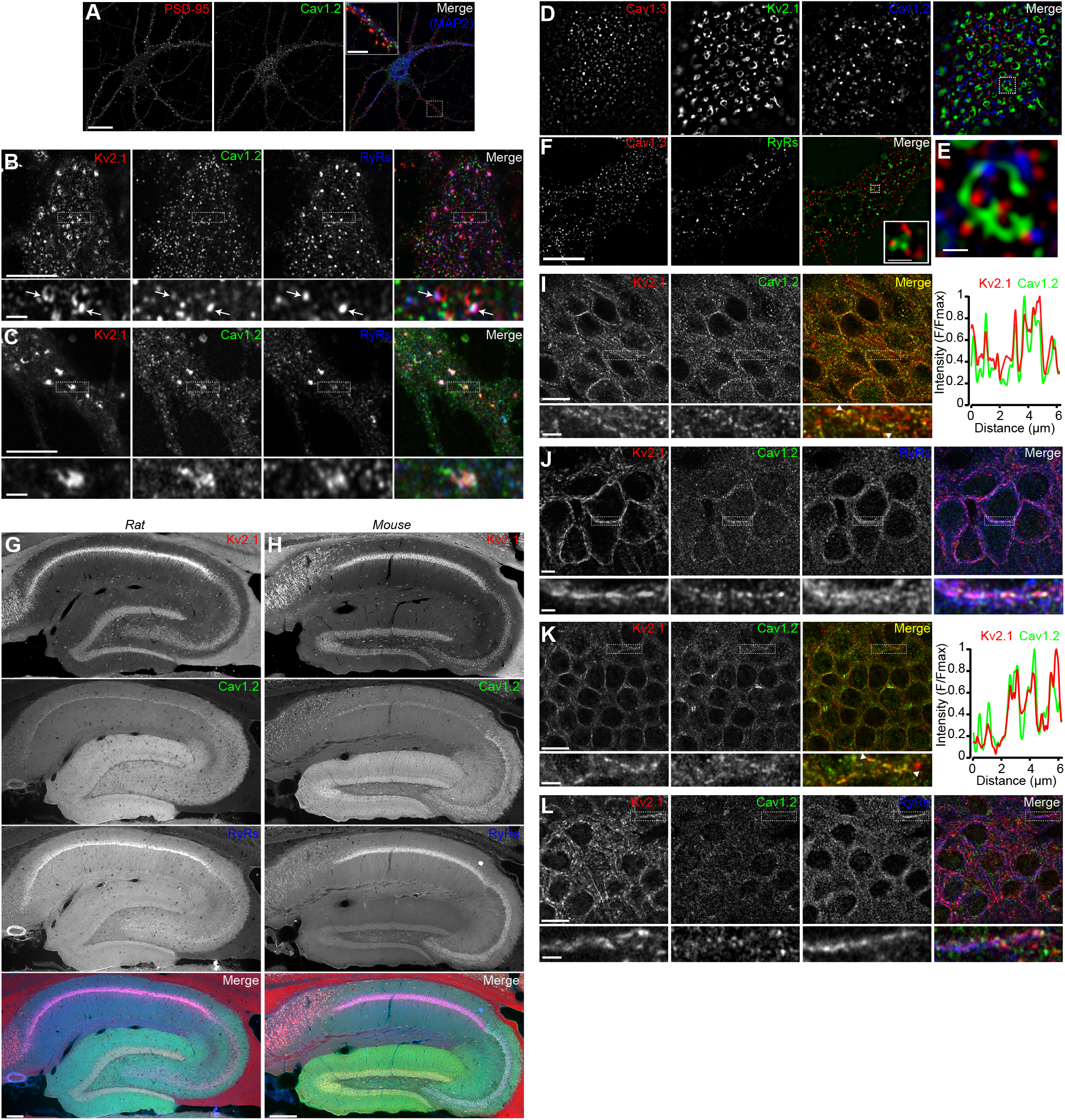
Kv2.1 spatially associates with Cav1.2 and RyRs in brain neurons. (A) Single optical section image of a rat CHN immunolabeled for PSD-95, Cav1.2, and MAP2 (scale bar: 20 μm). Note large population of somatic Cav1.2 channels distant from excitatory synapses located primarily on more distal dendrites. Inset of merged panel shows expanded view of dendritic PSD-95 and Cav1.2 immunolabeling marked by box (inset scale bar: 5 μm). (B) Single confocal optical section of the soma of rat CHN immunolabeled for Kv2.1, Cav1.2, and RyRs (scale bar: 5 μm). The row of panels below the main panels shows an expanded view of somatic immunolabeling in the region marked by the box in the main panels; arrows indicate selected regions of colocalized Kv2.1, Cav1.2, and RyR immunolabeling (inset scale bar: 1 μm). (C) As in B, but in a CHN displaying more prominent colocalization of clustered Kv2.1, Cav1.2, and RyRs. (D) Super resolution (N-SIM) image of the basal membrane of the soma of a rat CHN immunolabeled for Kv2.1, Cav1.2, and Cav1.3 (scale bar: 5 μm). (E) Expanded view of the boxed region in the merged image of D (scale bar: 1.25 μm). (F) Super resolution (N-SIM) image of the basal membrane of the soma of a rat CHN immunolabeled for Cav1.3 and RyRs (scale bar: 5 μm). Inset in merged panel shows a higher magnification view of the boxed area (inset scale bar: 0.625 μm). (G) Panels show exemplar images of the hippocampus acquired from a brain section from an adult rat immunolabeled for Kv2.1 (red), Cav1.2, (green) and RyRs (blue), and the merged image (scale bar: 200 μm). (H) As in G, but acquired from an adult mouse brain section. (I) Confocal optical section obtained from the dentate gyrus of a rat brain section immunolabeled for Kv2.1 (red) and Cav1.2 (green) (scale bar: 10 μm). The row below the main panels shows expanded views of immunolabeling in the region marked by the box in the main panels; arrowheads indicate region selected for intensity profile line scan (scale bar: 2 μm). Line scan obtained from inset is shown to the right. (J) Confocal optical section obtained from the pyramidal cell layer of hippocampal area CA1 in a rat brain section immunolabeled for Kv2.1 (red), Cav1.2 (green), and RyRs (blue) (scale bar: 10 μm). The row below the main panels shows expanded view of immunolabeling in the region marked by the box in the main panels (scale bar: 2 μm). (K) As in I but acquired from a mouse brain section. (L) As in J but acquired from a mouse brain section.

We next evaluated how phosphorylation-dependent dispersal of Kv2.1 clusters influenced the localization of somatic Cav1.2 and RyRs in rat CHNs. One stimulus that results in dispersal of Kv2.1 in CHNs is acute elevation in intracellular Ca^2+^ caused by the excitatory neurotransmitter glutamate (Misonou et al., 2004; Misonou et al., 2006). We found that glutamate stimulation of CHNs not only reduced Kv2.1 clustering, but also significantly decreased the colocalization between Cav1.2 and RyRs and increased the distance between somatic Cav1.2 clusters (Table 1). We also found that glutamate stimulation decreased the number of Cav1.2 clusters present on the PM, consistent with previous observations that acute Ca^2+^ influx results in endocytosis of Cav1.2 channels (Hall et al., 2013). Together, these data show that declustering of Kv2.1 in the PM is associated with reduced somatic coupling of Cav1.2 and RyR localization.

**Table 1.**
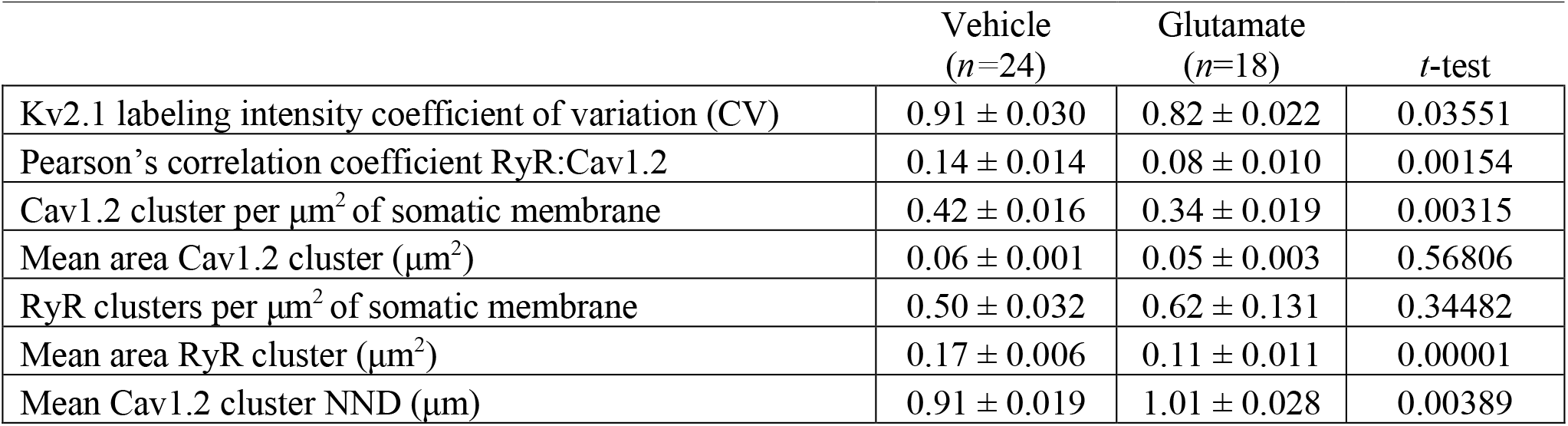
Cav1.2 and RyR colocalization parameters in rat CHNs.

We next assessed the localization of Kv2.1, Cav1.2, and RyRs in brain sections. Previous immunohistochemical analyses showed that in hippocampal neurons, Cav1.2 localizes to distinct clusters on somata and proximal dendrites (Westenbroek et al., 1990; Hell et al., 1993), a spatial pattern similar to that of Kv2.1 (Trimmer, 1991; Scannevin et al., 1996; Kirizs et al., 2014). Similar to previous observations, in low magnification images of mouse and rat hippocampus, we observed Cav1.2 immunolabeling concentrated in CA1 neuron somata, with increasing labeling in area CA2/CA3 neurons, and greatest labeling in dentate gyrus (DG) granule cell somata and dendrites (Fig. 1G-H). In higher magnification confocal images of DG granule cell bodies, we found that Kv2.1 clusters tended to colocalize with Cav1.2 clusters (Fig. 1I). The somata of CA1 pyramidal neurons had less intense Cav1.2 immunoreactivity, and colabeling with Kv2.1 was not as pronounced as in DG granule cells; however, the spatial association of Cav1.2, Kv2.1, and RyR immunolabeling in these cells was comparable to CHNs (Fig. 1J). Similar labeling was observed in high-magnification images of mouse brain sections (Fig. 1K-L). Kv2.2, which also clusters at EPJs through the same mechanism as Kv2.1 (Kirmiz et al., 2018b), similarly colocalized with Cav1.2 immunolabeling in CA1 pyramidal cells and DG granule cells (Fig. S1).

### Crosslinking-based proteomic analyses support that Kv2.1 channels are in close spatial proximity to LTCCs and RyRs in brain neurons

These findings indicated that LTCCs are spatially associated with Kv2.1 and RyRs in brain neurons. We next interrogated proteins within the Kv2.1 nano-environment using a crosslinking- and mass spectrometry-based proteomics approach to determine whether LTCCs and RyRs are in close spatial proximity (having lysine residues within ≈ 12 Å of one another) to Kv2.1. We affinity immunopurified (IPed) Kv2.1 from mouse brain homogenates that were subjected to chemical cross-linking during homogenization. This strategy previously allowed us to identify the ER-resident VAP proteins as Kv2 channel binding partners (Kirmiz et al., 2018b). Importantly, we also performed parallel IPs from brain homogenates prepared from Kv2.1 knockout (KO) mice (Jacobson et al., 2007; Speca et al., 2014) using the same Kv2.1 antibody, to identify proteins IPing in a Kv2.1-independent manner. To further improve the recovery of peptides IPed with Kv2.1, we performed on-bead trypsin digestion (Fig. S2), as opposed to the in-gel digestion we had done previously (Kirmiz et al., 2018b). Similar to our earlier findings, enriched in the control Kv2.1 IPs (and otherwise absent from the Kv2.1 KO brain IPs) were the VAP isoforms VAPA and VAPB (Table 1). In addition, among the most abundant 50 proteins specifically present in Kv2.1 IPs (*i.e.*, from WT and not Kv2.1 KO brain samples) were numerous proteins involved in Ca^2+^ signaling and/or previously reported to localize to neuronal EPJs. These included RyR isoforms RyR2 and RyR3, the LTCC α subunits Cav1.2 and Cav1.3, various Cavβ auxiliary subunits of LTCCs, as well as other proteins involved in Ca^2+^ signaling and homeostasis (Table 2). Taken together with our imaging analyses, these findings indicate that Kv2.1 is in close spatial proximity to LTCCs and RyRs at EPJs in mouse brain neurons. We note that while Cav1.2 is the predominant LTCC α1 subunit in hippocampus (Hell et al., 1993; Davare et al., 2001; Moosmang et al., 2005; Lacinova et al., 2008; Sinnegger-Brauns et al., 2009), where its localization on neuronal somata overlaps with Kv2.1, it was not as highly represented in these proteomic analyses as was Cav1.3, perhaps as these analyses were performed on whole brain samples.

**Table 2.** LTCC subunits and other Ca^2+^ signaling proteins specifically copurifying with Kv2.1.

| Protein | Rank | Mean | SEM (n=3) |
| --- | --- | --- | --- |
| Kv2.1 | 1 | 100.000 | NA |
| Kv2.2 | 3 | 31.638 | 0.518 |
| VAPA | 5 | 25.344 | 1.733 |
| RyR3 | 10 | 12.477 | 0.881 |
| Cavβ4 | 12 | 11.133 | 1.411 |
| VAPB | 15 | 7.600 | 1.393 |
| Cavβ2 | 18 | 5.623 | 0.79 |
| Cav1.3 | 19 | 5.730 | 1.652 |
| Cavβ3 | 23 | 5.070 | 1.033 |
| Hippocalcin | 24 | 4.583 | 0.831 |
| Neurocalcin-delta | 25 | 4.590 | 0.856 |
| SR/ER calcium ATPase 2 | 28 | 4.226 | 2.4 |
| Hippocalcin-like protein 1 | 29 | 4.360 | 0.288 |
| Cavβ1 | 33 | 3.800 | 0.697 |
| Calcineurin catalytic subunit γ | 35 | 3.583 | 0.718 |
| RyR2 | 36 | 3.140 | 0.903 |
| Calcineurin subunit B | 37 | 3.197 | 0.469 |
| Calcium-transporting ATPase | 39 | 2.873 | 0.447 |
| SR/ER calcium ATPase 1 | 40 | 2.530 | 1.21 |
| Cav1.2 | 43 | 2.427 | 0.766 |

### Kv2.1 organizes the localization of cell surface LTCCs

Because our immunolabeling and proteomics results indicated that endogenous Cav1.2 channels spatially associate with clustered Kv2.1 in hippocampal neurons, we next investigated how the subcellular localization of Cav1.2 (expressed with the LTCC auxiliary subunits α_2_δ_1_ and β3) was influenced by the presence of Kv2.1 in heterologous HEK293T cells. HEK293T cells lack endogenous Kv2.1 or Kv2.2 channels (Yu and Kerchner, 1998), and have little to no expression of LTCCs (Berjukow et al., 1996; Geiger et al., 2012). Expression of conducting or non-conducting Kv2 channels in these cells induces EPJ formation (Fox et al., 2015; Bishop et al., 2018; Kirmiz et al., 2018b). Using total internal reflection fluorescence (TIRF) microscopy to visualize Cav1.2-GFP expressed in HEK293T cells, we observed small (0.27±0.24 μm^2^) Cav1.2 clusters adjacent to cortical ER marked by the general ER marker BFP-SEC61b (Fig 2A). However, in the presence of Kv2.1, the PM organization of Cav1.2 was dramatically altered, such that Cav1.2 now co-assembled with Kv2.1 into significantly larger clusters (1.05±0.67 μm^2^) (Fig 2A-B). The Kv2.1-induced rearrangement of Cav1.2 was accompanied by an increased occurrence of larger Cav1.2 clusters and a reduced occurrence of smaller Cav1.2 clusters, and a nearly linear relationship between the sizes of Cav1.2 and Kv2.1 clusters (Fig 2B). Kv2.2 channels similarly recruited Cav1.2 into large clusters (Fig 2C). We determined that the impact of Kv2.1 expression on Cav1.2 clustering did not require Kv2.1 K^+^ conductance, as coexpression of a K^+^-impermeable point mutant (Kv2.1_P404W_) induced clustering of Cav1.2 comparable to WT Kv2.1 (Fig. 2D-E). Conversely, coexpression with a Kv2.1 point mutant (Kv2.1_S586A_), deficient in clustering (Lim et al., 2000) and in inducing EPJ formation (Kirmiz et al., 2018b), had no effect on Cav1.2 clustering (Fig. 2D-E). We also found that the localization of GFP-tagged Cav1.3 was similarly altered upon coexpression with Kv2.1 or Kv2.2, implying a common mechanism for co-clustering of LTCCs with Kv2 channels (Fig S3).

**Figure 2.**
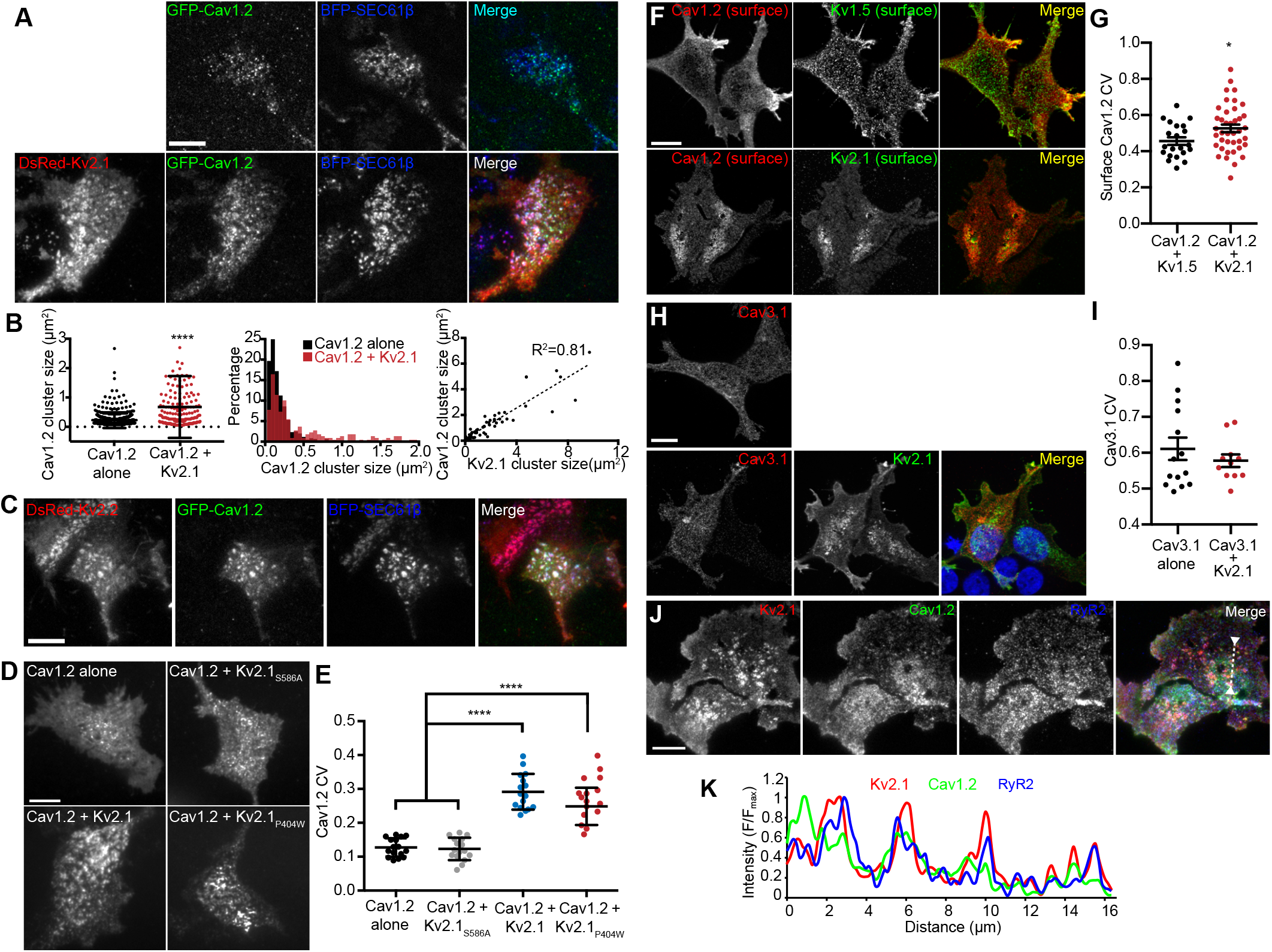
LTCCs are recruited to Kv2-induced EPJs. (A) Upper row: TIRF images of a HEK293T cell cotransfected with GFP-Cav1.2 (green), BFP-SEC61β (blue) and LTCC auxiliary subunits Cavβ3 and Cavα_2_δ_1_ (not shown) and without Kv2.1 (scale bar: 10 μm). Lower row: as in upper row, but in a cell additionally cotransfected with DsRed-Kv2.1. (B) Summary graphs of Cav1.2 cluster size (left panel), the cluster size frequency distribution (center panel), and a scatterplot of paired measurements of Kv2.1 and Cav1.2 cluster sizes (left panel) measured from HEK293T cells transfected with GFP-Cav1.2, Cavβ3, and Cavα_2_δ_1_ alone (black) or additionally cotransfected with DsRed-Kv2.1 (red) Bars are mean ±SD (****p <10^-15^, two-tailed *t*-test, *n*=3 cells). (C) TIRF images of a HEK293T cell cotransfected with DsRed-Kv2.2 (red), GFP-Cav1.2 (green), BFP-SEC61β (blue) and Cavβ3 and Cavα_2_δ_1_ (not shown). (D) TIRF images GFP-Cav1.2 in HEK293T cells cotransfected with GFP-Cav1.2, Cavβ3 and Cavα_2_δ_1_, either alone or with the non-clustered Kv2.1_S586A_ point mutant, Kv2.1_WT_, or the non-conducting Kv2.1_P404W_ point mutant (scale bar: 10 μm and holds for all panels). (E) Summary graph of coefficient of variation (CV) values of GFP-Cav1.2 fluorescent signal intensity measured from HEK293T cells cotransfected with GFP-Cav1.2 and the indicated Kv2.1 isoforms. Each point corresponds to a single cell. Bars are mean ±SD (Cav1.2 alone vs. Kv2.1_S586A_, p=0.6914; Cav1.2 alone vs. Kv2.1WT, ****p=3.904×10^-12^; Cav1.2 alone vs. Kv2.1_P404W_, ****p=7.812×10^-9^; two-tailed t-test). (F) Optical sections of HEK293T cells transfected with and immunolabeled for surface Cav1.2-HA and Kv1.5 (upper panels) or Kv2.1 (lower panels) (scale bar: 10 μm and holds for all panels). (G) Summary graph of CV values of Cav1.2-HA fluorescent signal intensity measured from HEK293T cells cotransfected with Kv1.5 or Kv2.1. Each point corresponds to a single cell (*p=0.0348 versus Kv1.5, two-tailed *t*-test). (H) Optical sections of HEK293T cells transfected with and immunolabeled for Cav3.1 alone (upper panel) or with Kv2.1 (lower panels) (scale bar: 10 μm and holds for all panels). (I) Summary graph of CV values of Cav3.1 fluorescent signal intensity measured from HEK293T cells described in H. Each point corresponds to a single cell (p=0.4027, two-tailed *t*-test). (J) TIRF images of a HEK293T cell cotransfected with DsRed-Kv2.1 (red), Cav1.2 (green), YFP-RyR2 (blue), and auxiliary subunits Cavβ3, Cavα_2_δ_1_, and STAC1 (not shown) (scale bar: 10 μm). (K) Line scan of fluorescence signal intensities of ROI depicted in J.

Because TIRF microscopy illuminates subcellular structures up to 100 nm away from the PM, we confirmed whether the observed co-clustering of Cav1.2 with Kv2.1 was occurring within the PM itself. We performed cell surface immunolabeling of intact cells coexpressing Kv2.1 and a Cav1.2 construct possessing an extracellular hemagglutinin epitope tag [Cav1.2-HA, (Obermair et al., 2004)]. Similar to cells expressing fluorescently tagged channels and imaged using TIRF microscopy, we found that PM-localized Cav1.2-HA co-clustered with PM Kv2.1, whereas the unrelated Kv1.5 channel did not cluster or associate with Cav1.2 (Fig. 2F-G). Moreover, Kv2.1 coexpression did not alter the PM localization of the T-type Ca^2+^ channel Cav3.1 (Fig. 2H-I). This observation suggests that the Kv2.1-mediated spatial reorganization of LTCCs is specific to their association with Kv2.1, a notion also supported by the absence of T-type Ca^2+^ channels from our Kv2.1 IP experiments.

We also assessed whether RyRs could be recruited to EPJs induced by Kv2.1. For these experiments, we expressed Kv2.1 along with Cav1.2, the LTCC auxiliary subunits α_2_δ_1_ and β3, RyR2, and the STAC1 adaptor protein, an approach similar to that previously used to recapitulate Cav1.1- and RyR1-mediated Ca^2+^ release in HEK293T cells (Perni et al., 2017). We found that in the presence of these auxiliary subunits, Kv2.1, Cav1.2, and RyR2 could spatially associate in HEK293T cells, similar to their association in hippocampal neurons (Fig. 2J-K). Together, these data demonstrate that clustered but not non-clustered Kv2 channels enhance LTCC clustering and increase their localization to EPJs as a nonconducting function, and that the spatial association of Kv2.1, Cav1.2, and RyRs seen in neurons can be recapitulated in HEK293T cells.

### Neuronal Kv2.1 channels functionally associate with endogenous LTCCs and RyRs

Kv2.1, when fused to fluorescent proteins such as GFP, clusters at neuronal EPJs similar to untagged or endogenous Kv2.1 (Antonucci et al., 2001; Kirmiz et al., 2018b). To begin to evaluate Ca^2+^ signals at neuronal Kv2.1-associated EPJs, we fused the genetically-encoded Ca^2+^ indicator GCaMP3 (derived from GFP) to K^+^-conducting and -nonconducting Kv2.1 channel isoforms and expressed these constructs in rat CHNs. GCaMP3 has previously been used to study near-membrane Ca^2+^ signaling microdomains in astrocytes (Shigetomi et al., 2010), and its higher basal fluorescence relative to newer GCaMP variants facilitated identification of transfected neurons. In rat CHNs, GCaMP3-Kv2.1 exhibited clustered localization similar to other fluorescently tagged Kv2.1 isoforms (Fig. 3A) and reported global Ca^2+^ spikes, as indicated by the synchronized increase in fluorescence across the PM at sites where the construct was clustered and also in regions with diffuse GCaMP3-Kv2.1 expression (Fig. 3B, movie S1). In addition to synchronized Ca^2+^ spikes, we also observed rapid and stochastic Ca^2+^ signals occurring at a subset of individual GCaMP3-Kv2.1 clusters within the soma (Fig. 3B-C, movie S1). These Ca^2+^ signals were confined to individual clusters such that the fluorescence of adjacent GCaMP3-Kv2.1 clusters <1 µm from the active clusters remained stable (Fig. 3B, compare regions of interest 2 and 4). We found that Ca^2+^ signal amplitude, frequency, and width were insensitive to the K^+^ conductance of the GCaMP3-Kv2.1 reporter, as Ca^2+^ signals detected by a K^+^-impermeable variant of this construct (GCaMP3-Kv2.1_P404W_) showed no difference in any of these parameters relative to GCaMP3-Kv2.1 (Fig. 3D).

**Figure 3.**
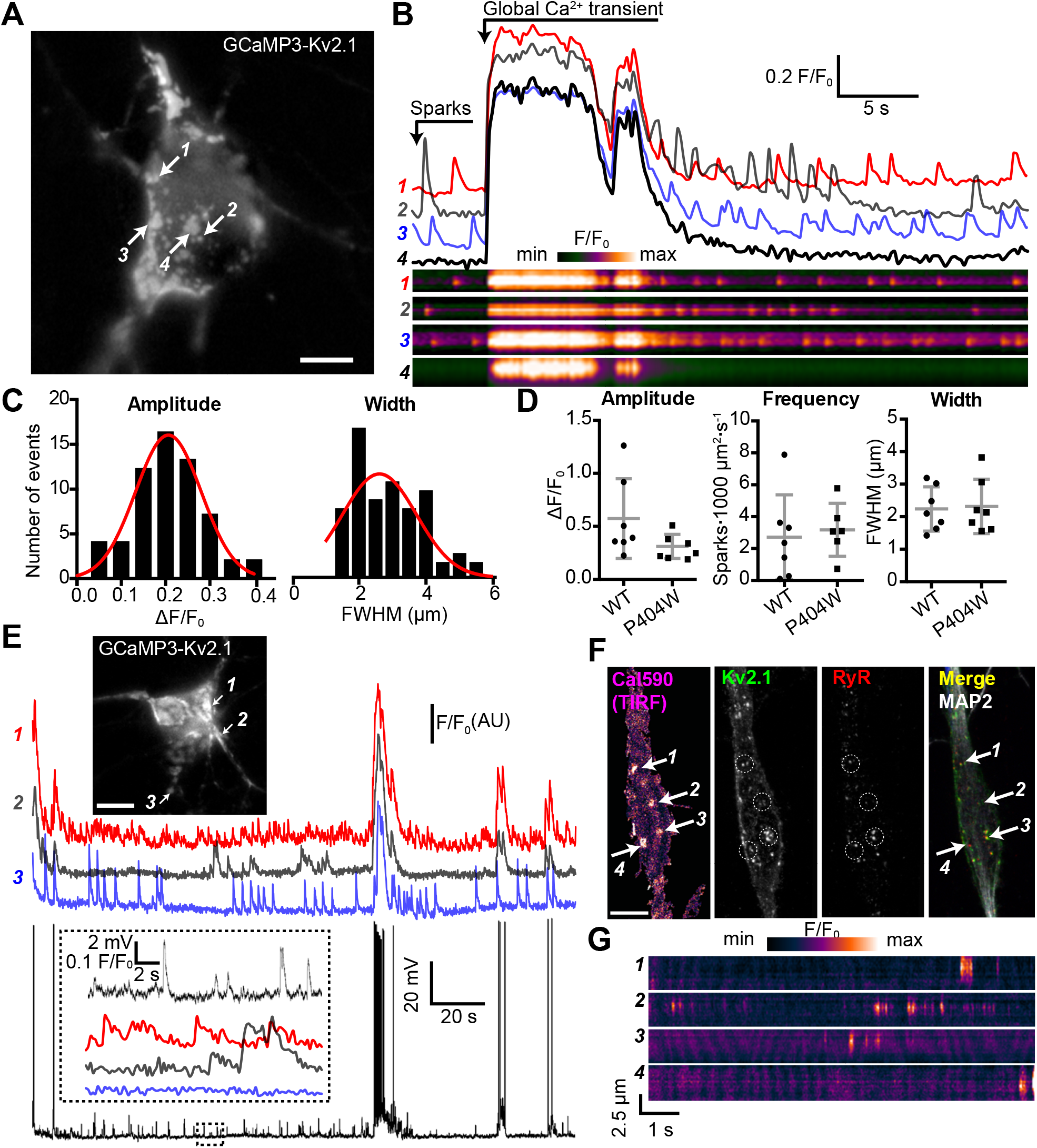
Spontaneous Ca^2+^ signals are generated at Kv2.1-associated EPJs. (A) Widefield image of a rat CHN transfected with GCaMP3-Kv2.1 (also see movie S1). Arrows indicate selected Kv2.1 clusters whose fluorescent intensity profiles are plotted in panel B (scale bar: 10 µm). (B) Fluorescence intensity traces (upper panels) and kymographs (lower panels) corresponding to the four ROIs indicated in panel A. Note spontaneous sparks occurring at ROI 2 that are not detected by the adjacent ROI 4. (C) Amplitude (ΔF/F_0_) and spatial spread (full width at half maximum, FWHM; µm) of all spatially distinct localized Ca^2+^ signals recorded from the neuron in panel A over a period of 90 seconds. (D) Summary data of the amplitude, frequency and spatial spread (width) of all spatially distinct localized Ca^2+^ signals recorded from CHNs expressing GCaMP3-Kv2.1 or GCaMP3-Kv2.1_P404W_. Each point corresponds to a single cell. No significant differences were detected. Bars are mean ±SD (Student’s t test). (E) Image of a rat CHN transfected with GCaMP3-Kv2.1 from which simultaneous GCaMP3-Kv2.1 fluorescence and membrane potential values were acquired (scale bar: 10 μm). Numbered arrows correspond to ROIs whose fluorescence intensity traces are depicted below image. Membrane potential measurements are provided in the bottom trace. The inset shows and expanded view of ROI Ca^2+^ traces and membrane potential values from region of the time course indicated by the dashed box in the membrane potential trace. (F) Representative rat CHN loaded with Cal590 and imaged with TIRF microscopy, followed by *post-hoc* immunolabeling for Kv2.1, RyRs, and MAP2. Arrows indicate ROIs where spontaneous Ca^2+^ signals were detected; dashed circles indicate approximate regions where immunolabeling for Kv2.1 and RyRs was detectable (scale bar: 10 μm). (G) Kymograph showing the localized Ca^2+^ release events detected at ROIs depicted in F.

Next, we assessed the relationship between GCaMP3-Kv2.1 reported Ca^2+^ signals and membrane potential (*V*_m_). We performed current clamp experiments to monitor the *V*_m_ and sparks simultaneously, using the whole-cell perforated patch clamp configuration. Spontaneous action potentials were associated with Ca^2+^ spikes, suggesting that these synchronized, large-amplitude Ca^2+^ transients reflected Ca^2+^ entry through voltage-gated Ca^2+^ channels as well as release through RyRs (Fig. 3E). However, unlike global Ca^2+^ spikes, the localized Ca^2+^ signals displayed no clear relationship with action potentials or other spontaneous *V*_m_ fluctuations, similar to previous observations of localized Ca2+ release events in CA1 pyramidal neurons (Berrout and Isokawa, 2009; Manita and Ross, 2009).

As heterologous expression of Kv2.1 in CHNs is known to result in large Kv2.1 “macroclusters” that recruit RyRs (Antonucci et al., 2001), we next determined whether somatic Ca^2+^ signals occurred at native Kv2.1-associated EPJs. For these experiments, we used non-transfected CHNs loaded with the Ca^2+^ dye Cal-590 AM and recorded sparks using TIRF microscopy. Using this approach, it was possible to detect spontaneous, localized Ca^2+^ release events in the soma that were qualitatively similar to those recorded with GCaMP3-Kv2.1 (Fig. 3F-G, movie S2). *Post-hoc* immunolabeling of these CHNs for Kv2.1, RyRs, and the neuron-specific cytoskeletal protein MAP2 indicated that the observed localized Ca^2+^ signals occurred primarily within the soma at sites of colocalized Kv2.1 and RyR clusters (Fig. 3F).

These observations suggested that the Ca^2+^ signals observed at neuronal Kv2.1-associated EPJs reflected RyR-generated Ca^2+^ sparks. To further assess this possibility, we imaged GCaMP3-Kv2.1-expressing CHNs treated with compounds that modulate LTCC- and RyR-mediated CICR. We found that caffeine, which sensitizes RyRs to cytosolic Ca^2+^, enhanced the frequency of localized Ca^2+^ sparks (Fig. 4A, B, movie S3). In contrast, depletion of ER Ca^2+^ stores with the sarco-/endo-plasmic reticulum Ca^2+^ ATPase (SERCA) inhibitor thapsigargin led to an elimination of Ca^2+^ sparks (Fig. 4A-B). The functional coupling of dendritic LTCCs and RyRs in hippocampal neurons has previously been demonstrated by the impact of dihydropyridine (DHP) compounds on dendritic sparks: the LTCC agonist Bay K8644 increased spark frequency, whereas the LTCC inhibitor nimodipine blocked sparks (Manita and Ross, 2009). Here, we obtained similar evidence of the involvement of LTCCs in the generation of somatic GCaMP3-Kv2.1 reported Ca^2+^ sparks, whose frequency was enhanced by activation of LTCCs with Bay K8644 (Fig 4A, B, D, movie S4). Conversely, Ca^2+^ sparks were rapidly inhibited by blockade of LTCCs with nimodipine (Fig. 4A-B). We also performed *post-hoc* immunolabeling of imaged CHNs to determine whether the specific GCaMP3-Kv2.1 clusters which exhibited localized Ca^2+^ signals were associated with RyRs. Using this approach, we determined that the subset of GCaMP3-Kv2.1 clusters that colocalized with RyRs corresponded to the clusters that produced localized Ca^2+^ signals, either spontaneously or in response to the pharmacological modulators caffeine (Fig. 4C) and Bay K8644 (Fig. 4D). We also quantified the relationship between the size of *post-hoc* immunolabeled RyR clusters and spark frequency and amplitude. Similar to previous observations in vascular smooth muscle (Pritchard et al., 2018) and cardiac muscle (Galice et al., 2018) cells, we found that neuronal Ca^2+^ spark frequency but not amplitude correlated with RyR cluster size, and that application of the LTCC agonist Bay K8644 steepened this relationship (Fig. 4E). Taken together, these observations demonstrate that Kv2.1-associated EPJs are sites of spontaneous CICR events mediated by LTCCs and RyRs.

**Figure 4.**
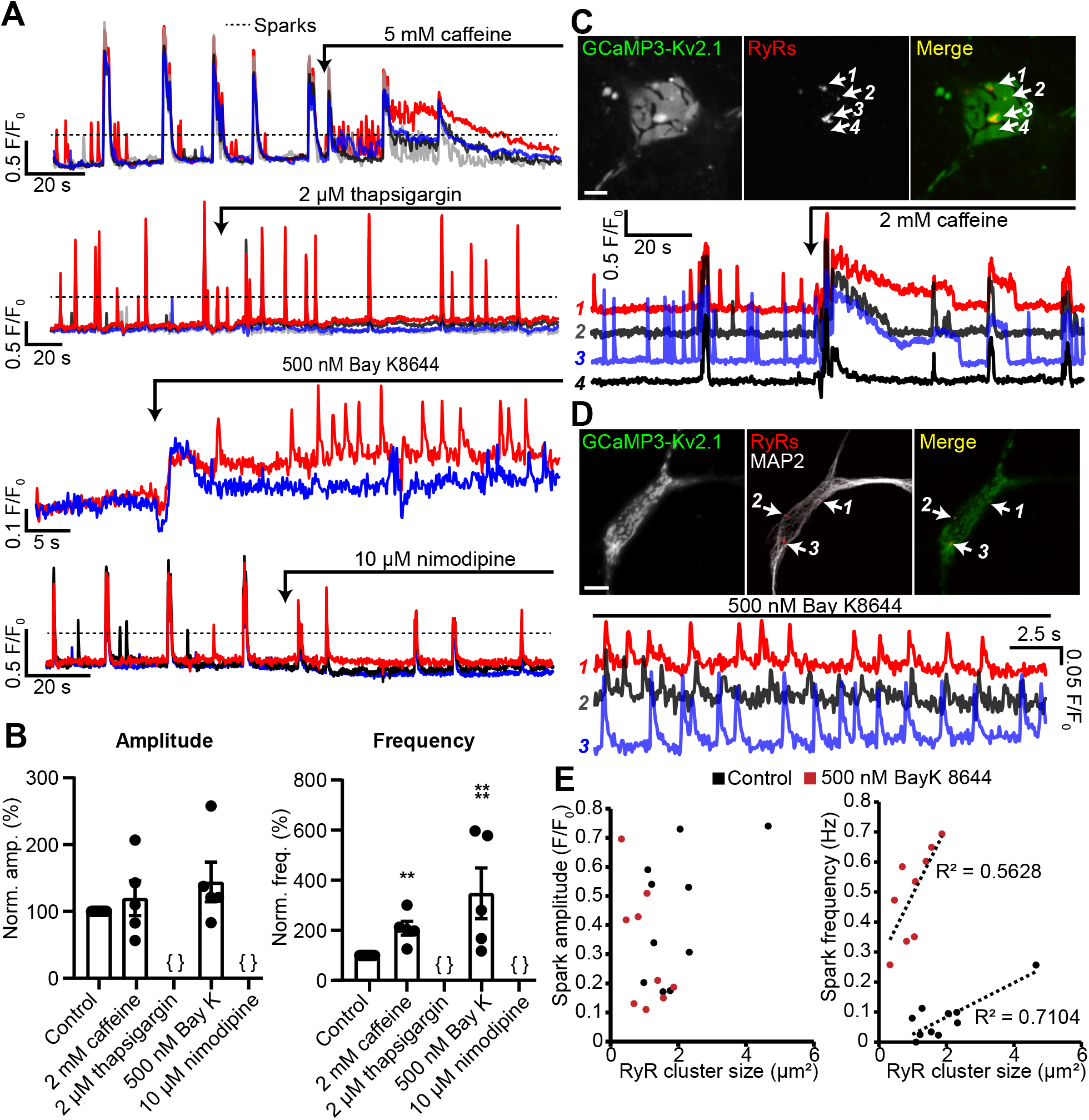
Spontaneous Ca^2+^ signals at Kv2.1-associated EPJs are produced by RyR- and LTCC-mediated CICR. (A) Representative GCaMP3-Kv2.1 fluorescence traces from CHNs treated with pharmacological probes of CICR. Different colors indicate spatially distinct ROIs within the same neuron. Dashed line indicates typical threshold for localized Ca^2+^ signals as opposed to the larger amplitude, synchronized global Ca^2+^ transients. (B) Summary data of the amplitude and frequency of all sparks recorded from CHNs treated with pharmacological probes of CICR. Each point corresponds to a single cell (**p=0.0013 vs. control; ****p<0.0001 vs. control; { }: no Ca^2+^ sparks detected; One-way ANOVA followed by Dunnett’s test). (C) Image of rat CHN transfected with GCaMP3-Kv2.1 and treated with caffeine, followed by *post-hoc* immunolabeling for RyRs (scale bar: 10 μm). Numbered arrows indicate ROIs where localized Ca^2+^ signals were detected (ROIs 1-3) or not detected (ROI 4). ROI fluorescence traces are shown in lower panel; note lack of spontaneous Ca^2+^ signals at ROI 4 despite its proximity to ROI 3, which displays prominent spontaneous Ca^2+^ release. (D) As in panel A, except CHN was treated with 500 nM Bay K8644 to induce spontaneous Ca^2+^ signals (scale bar: 10 μm). (E) Plot of individual RyR cluster size (determined from *post-hoc* immunolabeling) versus its spark amplitude (left panel) or frequency (right panel) reported by GCaMP3-Kv2.1 fluorescence in control (black symbols) or Bay K8644-treated (red symbols) cells. Each point corresponds to an individual RyR cluster (*n*=data from 4 cells [control] or 5 cells [Bay K8644]).

### Kv2.1 augments LTCC and RyR2-mediated CICR reconstituted in HEK293T cells

We next asked how Kv2.1-induced clustering of LTCCs would impact RyR-mediated Ca^2+^ release in HEK293T cells. For these experiments, we expressed Kv2.1 along with Cav1.2, the LTCC auxiliary subunits α_2_δ_1_ and β3, RyR2, and the STAC1 adaptor protein, which leads to co-clustering of Kv2.1, Cav1.2 and RyR2, as demonstrated in Fig. 2J-K. To detect Ca^2+^ release events, we performed TIRF microscopy of cells loaded with the Ca^2+^-sensitive dye Cal-590 AM. Although it was not possible to establish whether a cell expressed all transfected constructs, we observed spontaneous Ca^2+^ release events in a subset of cells (Fig. 5A, D) that were not seen in untransfected HEK293T cells and focused our analysis on cells that exhibited this phenotype. These spontaneous Ca^2+^ release events were rapidly blocked by the RyR inhibitor tetracaine (Fig. 5F, movie S5), suggesting that they reflected CICR mediated by RyRs. Expressing Kv2.1 in these cells resulted in enhanced spark frequency and amplitude (Fig. 5B-C, E). Similar results were obtained using Cav1.3 in place of Cav1.2 (Fig. S4). To better understand the mechanism underlying the influence of Kv2.1 on these reconstituted Ca^2+^ sparks, we next compared how they were affected by the non-conducting Kv2.1_P404W_ and the non-clustering Kv2.1_S586A_ point mutants (Fig. 5H). By using these Kv2.1 isoforms, we determined that there was an interplay between both Kv2.1 K^+^ conductance and clustering on Ca^2+^ sparks reconstituted in HEK293T cells. Expression of Kv2.1 channels capable of clustered EPJ formation (i.e., Kv2.1_WT_ and Kv2.1_P404W_) increased spark frequency, whereas non-clustering Kv2.1_S586A_ did not (Fig. 5I). Interestingly, we found that spark amplitude was enhanced by K^+^-conducting Kv2.1_WT_ but not Kv2.1_P404W_, suggesting that while Kv2.1-mediated clustering alone was sufficient to impact spark frequency, K^+^ conductance was required to impact the amplitude of reconstituted Ca^2+^ sparks. In conclusion, these observations indicate that Kv2.1-mediated clustering promotes the functional coupling of Cav1.2 and RyRs.

**Figure 5.**
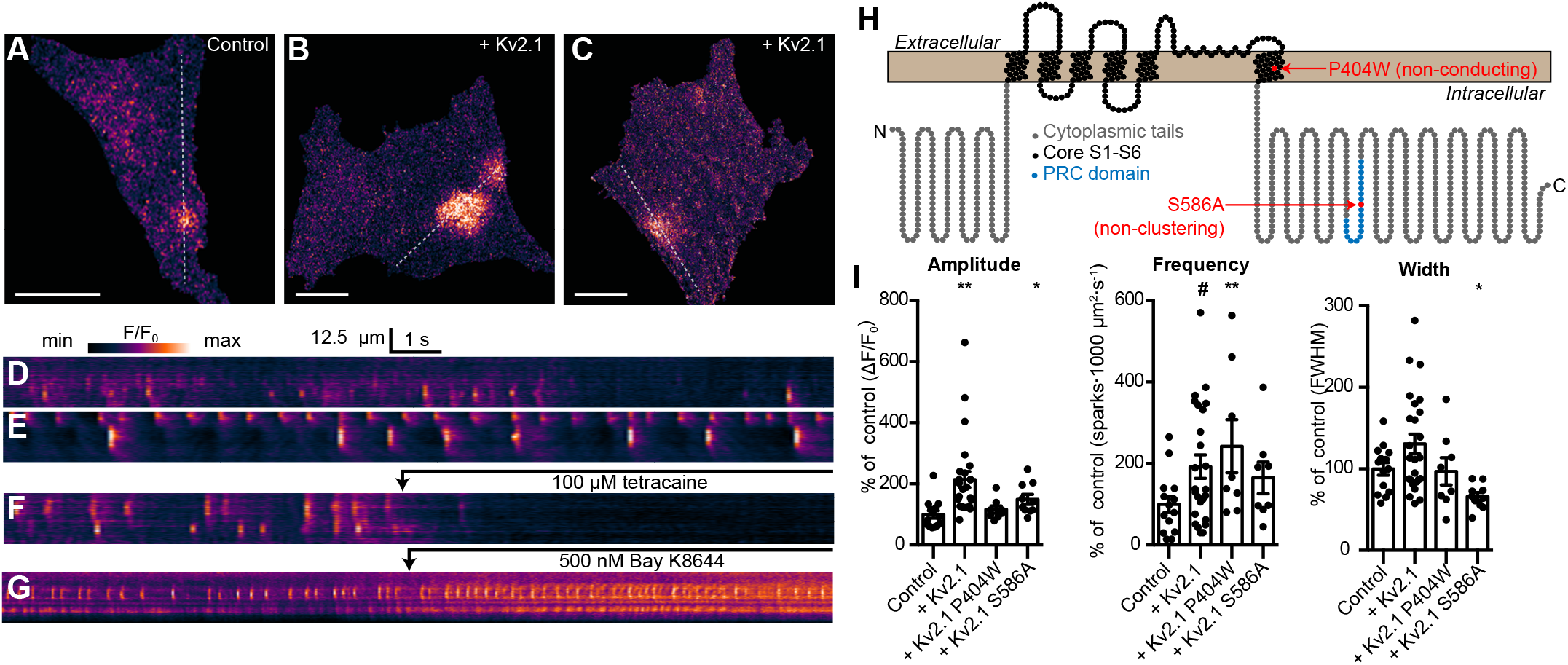
Kv2.1 expression increases the frequency of LTCC- and RyR-mediated sparks reconstituted in HEK293T cells. (A) TIRF image of a HEK293T cell expressing Cav1.2, RyR2, STAC1, and the LTCC auxiliary subunits β3 and α2δ1, and loaded with Cal-590 AM. (B-C) TIRF images of HEK293T cells additionally coexpressing Kv2.1. Dashed line indicates ROI depicted in corresponding kymographs (scale bar in panels A-C: 10 µm). (D-F) Kymograph showing the localized Ca^2+^ release events detected in the ROI on the cell in panels A-C, respectively. In (F), 100 µM tetracaine was added at the indicated time point. (G) Kymograph showing the localized Ca^2+^ release events detected in a cell treated with 500 nM Bay K8644 at the indicated time point. (H) Illustration of the membrane topology of a single Kv2.1 α subunit depicting the locations of the P404W and S586A point mutations. (I) Summary data of the amplitude, frequency and spatial spread (width) of all sparks recorded from HEK293T cells expressing Cav1.2, RyR2, and auxiliary subunits, without (control) or with addition of the indicated Kv2.1 isoforms. Each point corresponds to a single cell (width: *p=0.048; amplitude: **p<0.0001, *p=0.039; frequency: #p=0.053, **p=0.033; One-way ANOVA followed by Dunnett’s *post-hoc* test vs. control).

### Kv2.1 reduces the voltage threshold for Cav1.2 opening

Having demonstrated a spatial and functional association of Kv2.1, LTCCs, and RyRs in hippocampal neurons that could be reconstituted in HEK293T cells, we next investigated whether clustering by Kv2.1 influenced the Cav1.2-mediated LTCC activity. As physical interactions between adjacent LTCCs promote enhanced LTCC activity (reducing the membrane voltage threshold for channel opening and elevating channel open probability) (Navedo et al., 2005; Dixon et al., 2012; Moreno et al., 2016), we reasoned that this functional property of Cav1.2 might be enhanced by Kv2.1-induced clustering. To test this possibility, we obtained whole-cell patch-clamp recordings from HEK293T cells transfected with Cav1.2 and the non-K+ conducting Kv2.1_P404W_ point mutant, which allowed us to measure Ca^2+^ currents (*I*_Ca_) in the absence of the very large outward K^+^ currents produced by Kv2.1_WT_. Consistent with an influence of Cav1.2 spatial organization on its activity, we found that expression of Cav1.2 with Kv2.1_P404W_ more than doubled peak *I*_Ca_ as compared to cells expressing Cav1.2 alone (Fig. 6A-B). Analysis of the conductance-voltage (*G-V*) relationship also showed an influence of Kv2.1 on the *V*_m_ threshold for Cav1.2 opening, with currents produced by Cav1.2 activating at more negative voltages in the presence of Kv2.1_P404W_ than those produced by Cav1.2 alone, with no effect on steady-state inactivation (Fig. 6C). Cells co-expressing STAC1 with Cav1.2 and Kv2.1_P404W_ also exhibited an increase in whole-cell *I*_Ca_ and a hyperpolarized shift in Cav1.2 opening, similar to results obtained without STAC1 (Fig. S5). Measurement of Ca^2+^-induced fluorescence increases in cells loaded with the Ca^2+^-sensitive dye Rhod-2 via the patch pipette also revealed an enhancing effect of Kv2.1_P404W_ on Cav1.2-mediated Ca^2+^ influx (Fig. 6F). Similarly, HEK293T cells loaded with the Ca^2+^ dye Fluo-4 and expressing Cav1.2 and either Kv2.1_WT_ or Kv2.1_P404W_ displayed greater K^+^-depolarization induced Ca^2+^ influx than control cells (Fig. 6G-H), further supporting that K^+^-conducting as well as -nonconducting isoforms of Kv2.1 augment Cav1.2 activity.

**Figure 6.**
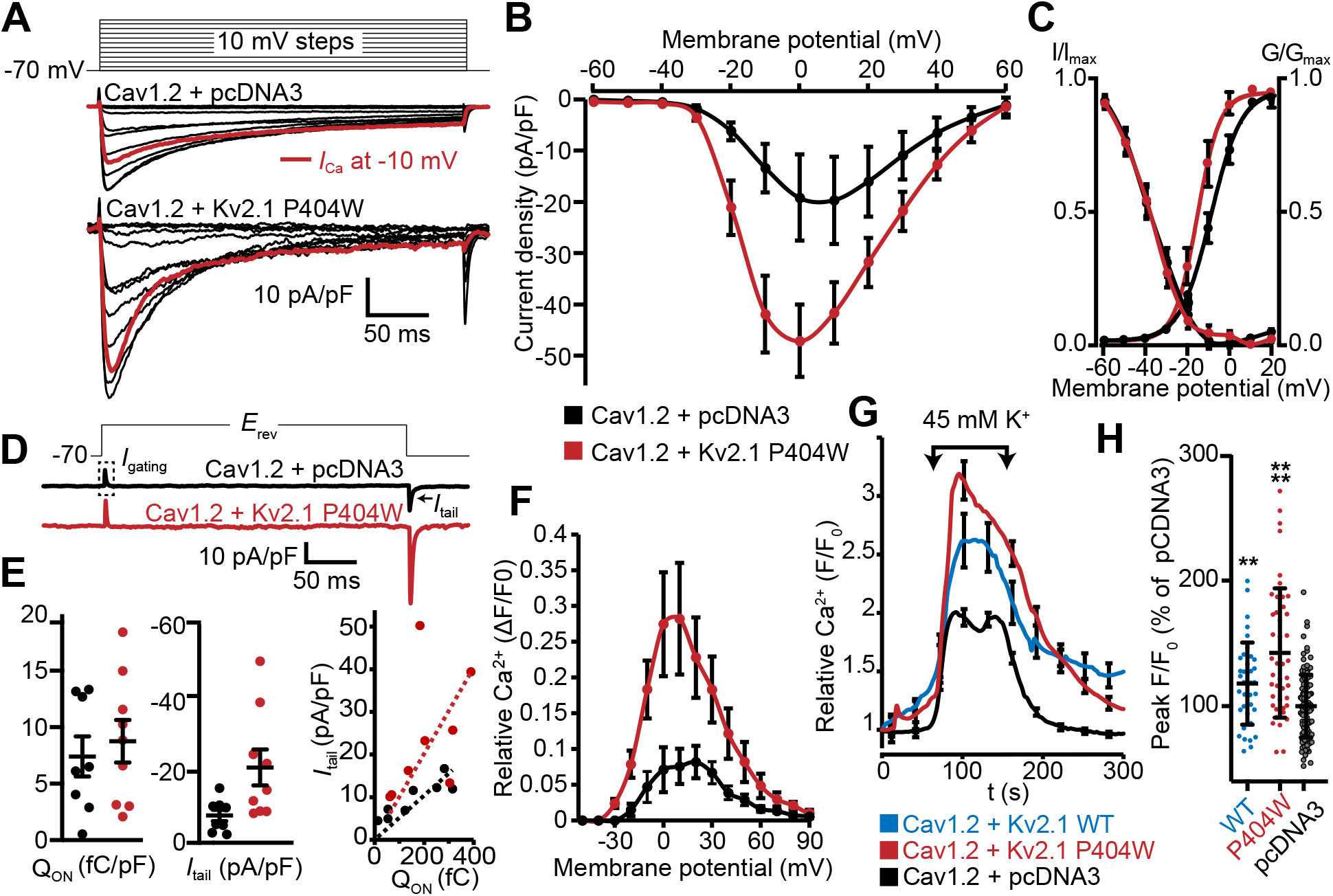
Cav1.2 channel activity is increased by coexpression with Kv2.1_P404W_. (A) Representative Ca^2+^ current trace families recorded from HEK293T cells transfected with Cav1.2-GFP and auxiliary subunits Cavβ3 and Cavα_2_δ_1_, without (+ pcDNA3 empty vector) with cotransfection of DsRed-Kv2.1_P404W_. For panels D-F, data are from cells without (+ pcDNA3 empty vector, in black) or with coexpression of Kv2.1_P404W_ (in red). (B) Normalized current-voltage (*I-V*) relationship of whole-cell *I*_Ca_ recorded from n=17 (Cav1.2 + pcDNA3) and *n*=10 (Cav1.2 + Kv2.1_P404W_) cells. (C) Voltage-dependence of whole cell Cav1.2 conductance *G*/*G*_max_ and steady-state inactivation *I*/*I*_max_. For the conductance-voltage relationships, the half-maximal activation voltage *V*_1/2_=-8.9±0.8 [pcDNA3] vs. -13.9±1.6 [+Kv2.1_P404W_] mV, p=0.0045; slope factor *k*=6.9±0.3 [pcDNA3] vs. 4.5±0.7 [+Kv2.1_P404W_], p=0.0025; Student’s t-test. (D) Representative nitrendipine-sensitive Cav1.2 gating and tail currents recorded from control (pcDNA3) cells and cells coexpressing Kv2.1_P404W_. (E) Quantification of nitrendipine-sensitive Cav1.2 Q_on_ (left), *I*_tail_ (center), and Q_on_ vs. *I*_tail_ (right).. Each point corresponds to a single cell (*p=0.019, Student’s *t*-test). (F) Average Rhod-2 fluorescence intensity measurements obtained from cells held at different membrane potentials during voltage clamp experiments (*n*=4 cells per condition). (G) Average fluorescence intensity measurements from Fluo4-loaded HEK293T cells transfected with Cav1.2, auxiliary subunits Cavβ3 and Cavα2δ, without (+ pcDNA3 empty vector, in black) or with Cotransfection of Kv2.1_WT_ (in blue) or Kv2.1_P404W_ (in red). Ca^2+^ influx was stimulated by depolarization with high extracellular K^+^ (45 mM) as indicated on the graph. (H) Average peak fluorescence values obtained during high-K^+^ depolarization of HEK293T cells expressing Cav1.2 and Kv2.1_WT_ or Kv2.1_P404W_ as in G. Each point corresponds to a single cell. Bars are mean ± SD (**p<0.0001, *p=0.0047 versus control; Student’s *t*-test).

Ion channel activity can be described by the product of the number of channels present in the PM (*n*), the channel’s unitary conductance (*i*), and the open probability of these channels (*P*_o_), such that the whole cell current *I* can be described by the relationship *I*=*nP*_o_*i*. Thus, the enhancement of Cav1.2 activity observed in the presence of Kv2.1 could be caused by an effect on any one or more of these parameters. To better understand the underlying mechanism, we acquired gating and ionic tail currents from the same cell. Depolarization-induced voltage sensor movement in activating voltage-gated channels produces a gating current (Q_on_) that is proportional to the number of channels present in the PM (*n*). Repolarization-induced ionic tail currents (*I*_tail_) reveal overall channel activity (*I*). Changes in one or both can be used to infer whether it is “*n*” versus some combination of “*P*_o_” and/or “*i*” that yield changes in total channel activity. We used nitrendipine, a DHP LTCC gating inhibitor, to pharmacologically isolate Cav1.2 Q_on_ when the *V*_m_ was stepped to the *I*_Ca_ reversal potential, and to measure *I*_tail_ elicited by returning to the -70 mV holding potential (Fig. 6D). Nitrendipine-sensitive Q_on_ values produced by Cav1.2 alone were comparable to those measured in the presence of Kv2.1, indicating that the increased *I*_Ca_ in cells coexpressing Kv2.1 was not associated with an increase in the number of PM Cav1.2 channels (Fig. 6E). However, the nitrendipine-sensitive *I*_tail_ was significantly greater in the presence of Kv2.1, demonstrating that the open probability and/or conductance of Cav1.2 was increased when co-expressed with Kv2.1. As comparable Q_on_ values (i.e., Cav1.2 voltage sensor movement) produced a larger *I*_tail_ in the presence of Kv2.1, taken together with the altered G-V curve shown in Fig. 6C suggests that the Kv2.1-dependent increase in *I*_Ca_ apparently came from enhanced Cav1.2 voltage sensor coupling to channel opening.

These observations showed that Cav1.2 channel activity was enhanced in the presence of Kv2.1. Therefore, we next asked whether LTCC currents were altered in CHNs lacking Kv2.1. For these experiments, we chose to record from CHNs as opposed to acutely dissociated neurons. Although the round morphology of acutely dissociated neurons enables much better control of the *V*_m_ than in arborized neurons, we reasoned based on the loss of Kv2.1 clustering upon dissociation in other cell types expressing clustered Kv2.1 (PC12, MDCK, and HEK293 cells; J.S. Trimmer, unpublished observations), and that endogenous Kv2.1 clusters in CHNs are sensitive to changes in intracellular Ca^2+^ and metabolism (Misonou et al., 2005a), that acute dissociation would disrupt the clustered localization of Kv2.1, potentially concealing LTCC regulation by Kv2.1 clustering. To improve somatic voltage clamp, we used recording solutions lacking Na^+^ and containing Cs^+^ and Ba^2+^ (which block K^+^ channels; Ba^2+^ also permeates voltage-gated Ca^2+^ channels) to increase membrane impedance. We focused our analyses of electrophysiological recordings on repolarization-induced tail currents after activation of channels by a depolarizing prepulse, rather than measurement of currents induced by depolarizing voltage steps that can be distorted due to space clamp limitations (e.g., see (Milescu et al., 2010). Similar to our findings in HEK293T cells, whole cell Ba^2+^ currents (*I*_Ba_) at +10 mV, as well as LTCC tail currents (Fig. 7B, C) were larger in CHNs from WT mice than those measured in Kv2.1 KO CHNs (Fig. 7A-C). To isolate the LTCC component of *I*_Ba_, we applied the LTCC gating inhibitor nimodipine (10 µM), and found that the reduced *I*_Ba_ observed in Kv2.1 KO CHNs (Fig. 7A-C) was primarily due to a reduction in the nimodipine-sensitive component of the current (Fig. 7A, B, E), with no apparent difference in the nimodipine-resistant current (Fig. 7A, B, D). We also examined nimodipine-sensitive gating and ionic tail currents when the *V*_m_ was stepped to the *I*_Ba_ reversal potential and found that while Q_on_ was not significantly different between control and Kv2.1 KO CHNs, peak Itail was reduced in Kv2.1 KO CHNs (Fig. 7F-G). The data in Fig. 6 (from exogenously expressed channels in HEK293T cells) and Fig. 7 (from endogenously expressed channels in CHNs) show that Kv2.1 enhances neuronal LTCC activity and suggest that the underlying mechanism involves enhanced coupling efficiency between LTCC voltage sensor movement and channel opening due to Kv2.1-mediated clustering.

**Figure 7.**
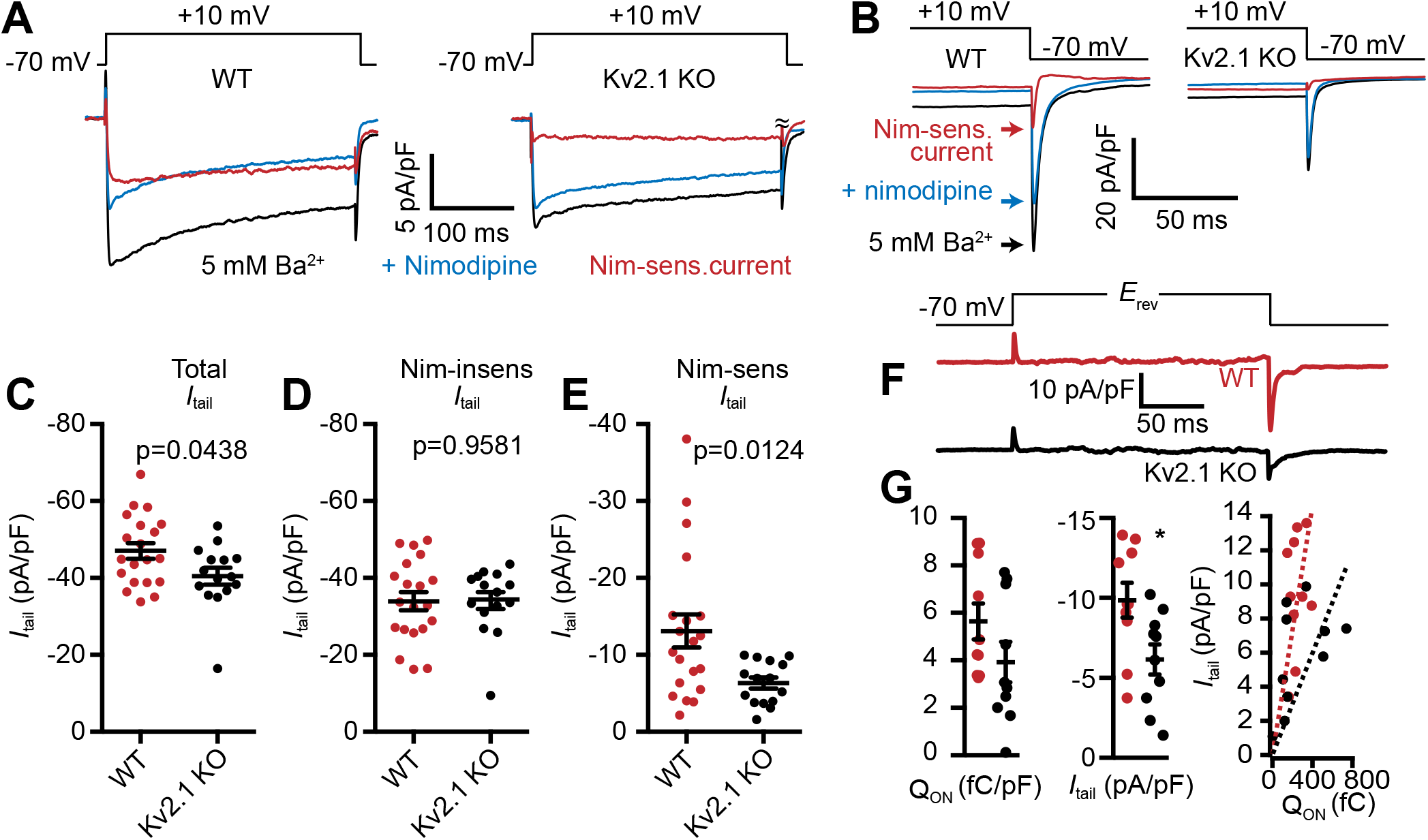
LTCC activity is reduced in Kv2.1 KO hippocampal neurons. (A) Representative Ba^2+^ current traces recorded from WT (left) and Kv2.1 KO CHNs (right) recorded at +10 mV in vehicle or in the presence of the LTCC inhibitor nimodipine (10 µM). (B) Representative raw tail current records from a WT (left) and Kv2.1 KO (right) CHN induced by a step to -70 mV from a 10 mV prepulse, recorded in vehicle or in the presence of 10 µM nimodipine. C-F. Comparison of WT (red) and Kv2.1 KO (black) CHNs. (C) Maximum tail current amplitudes measured at -70 mV from a 10 mV prepulse. Each point represents one cell. (D) As in C but recorded in the presence of 10 µM nimodipine. (E) Maximum nimodipine-sensitive tail current amplitudes obtained from each cell by subtracting maximum tail current amplitudes measured in vehicle from those measured in the presence of nimodipine. (F) Representative nimodipine-sensitive LTCC gating and tail currents recorded from WT and Kv2.1 KO CHNs. (G) Quantification of nimodipine-sensitive LTCC Q_on_ (left), *I*_tail_ (center), and Q_on_ vs. *I*_tail_ (right) recorded from WT and Kv2.1 KO CHNs. Each point corresponds to a single cell (*p=0.019, Student’s *t*-test).

### Kv2.1 promotes spatial coupling of LTCCs and RyRs

Given that Kv2.1-mediated clustering impacts the spatial distribution of Cav1.2 in coexpressing HEK293T cells, we next examined whether loss of Kv2.1 was associated with changes in the expression and localization of Cav1.2. We first performed immunolabeling of hippocampal neurons in brain sections from adult control and Kv2.1 KO mouse littermates. We have previously determined that the anatomic structure of mouse brains lacking Kv2.1 is comparable to controls, and there do not appear to be compensatory changes in the expression of other Kv channels tested (Speca et al., 2014). Here, we confirmed that immunolabeling for somatodendritic Kv2.2 and also dendritic Kv4.2 channels was similar in WT and Kv2.1 KO hippocampus (Fig. 8A-C). However, Cav1.2 labeling was increased in pyramidal neurons in area CA1 in Kv2.1 KO brain sections, both within the cell bodies and in the apical dendrites (Fig. 8C). These results suggest that in adult mice lacking functional Kv2.1 channels, Cav1.2 expression may be elevated, potentially as a compensatory mechanism to overcome reduced Cav1.2 channel function.

**Figure 8.**
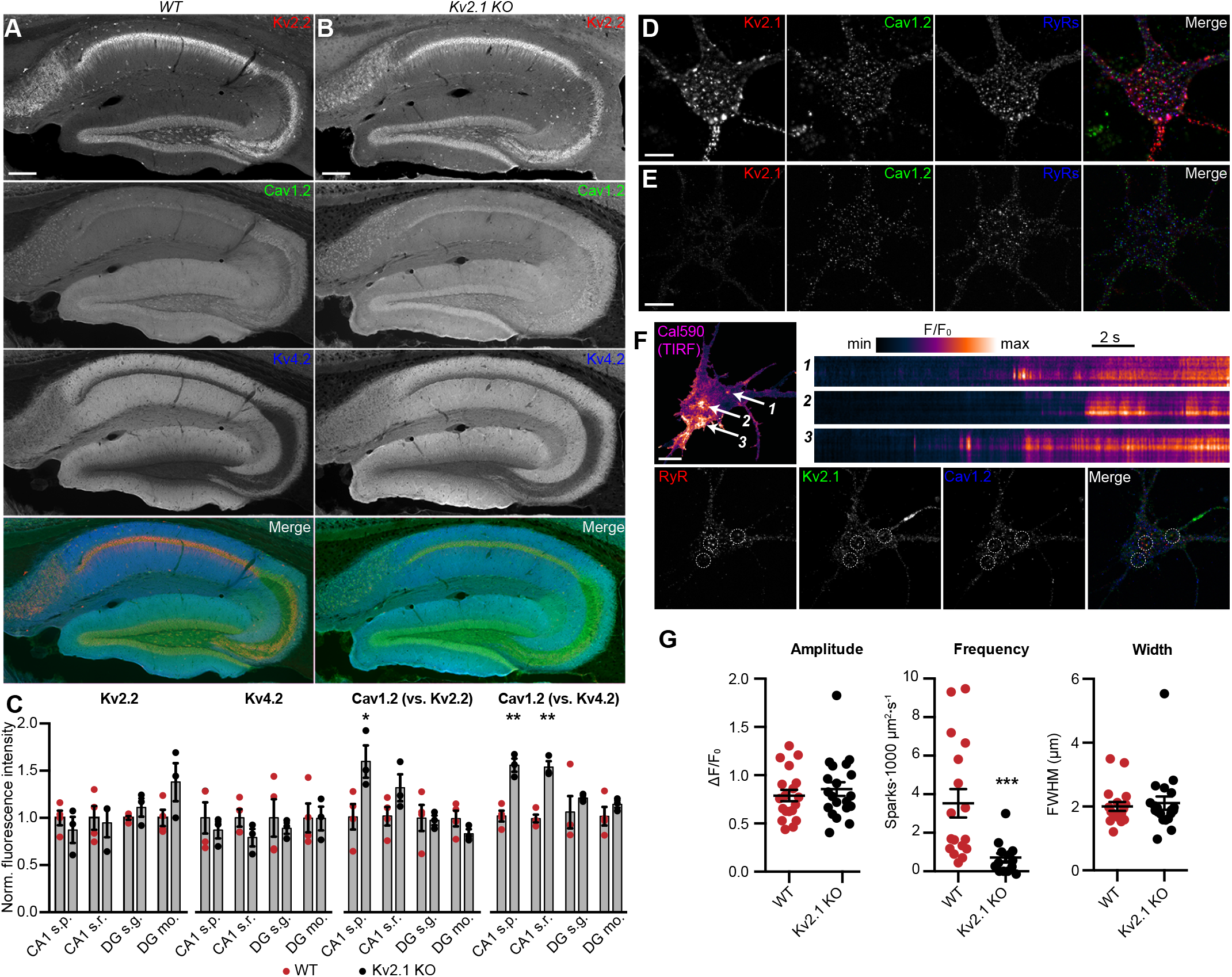
Increased immunolabeling for Cav1.2 in Kv2.1 KO brain sections, and reduced spark frequency in cultured Kv2.1 KO CHNs. (A) Column shows exemplar images of the hippocampus acquired from brain sections of adult WT mice immunolabeled for Kv2.2 (red), Cav1.2 (green) and Kv4.2 (blue) (scale bar: 200 μm). (B) As in A but acquired from Kv2.1 KO mice. (C) Summary graphs of normalized mean fluorescence intensity of Kv2.2, Kv4.2, and Cav1.2 immunolabeling from ROIs from various laminae within CA1 (s.p.: *stratum pyramidale*; s.r.: *stratum radiatum*) and DG (s.g.: stratum granulosum; mo: molecular layer) in brain sections from adult WT (red) and Kv2.1 KO (black) mice. Each point corresponds to an individual mouse (Cav1.2 vs. Kv2.2: *p=0.0408; Cav1.2 vs. Kv4.2: **p=0.0018, ***p=0.0007). (D) A single optical section image of a WT mouse CHN immunolabeled for Kv2.1, Cav1.2, and RyRs (scale bar: 10 μm). (E) As in D but acquired from a Kv2.1 KO mouse CHN. (F) Representative WT mouse CHN loaded with Cal590 and imaged with TIRF microscopy, followed by *post-hoc* immunolabeling for RyRs, Kv2.1, and Cav1.2. Arrows indicate ROIs where spontaneous Ca^2+^ signals were detected; dashed circles indicate approximate regions where immunolabeling for Kv2.1, Cav1.2, and RyRs was detectable. Kymograph showing the localized Ca^2+^ release events detected at ROIs are depicted to the right. (G) Summary data of the amplitude, frequency and spatial spread (width) of all sparks recorded from WT and Kv2.1 KO mouse CHNs. Each point corresponds to a single cell (***p=0.0042 versus WT; Student’s *t*-test).

To obtain more detailed individual cell information, we next investigated how the loss of endogenous Kv2.1 influenced the localization and function of LTCCs and RyRs in WT and Kv2.1 KO CHNs. To determine whether Kv2.1 channels regulate the localization of somatodendritic Cav1.2 and/or RyRs, we first analyzed the size and morphology of immunolabeled Cav1.2 and RyR clusters in control and Kv2.1 KO mouse CHNs. We found reduced colocalization between Cav1.2 clusters and RyR clusters in the absence of Kv2.1, and increased distance between Cav1.2 clusters (Fig. 8D-E, Table 3). In addition, we found that although the total number of somatic RyR clusters was not altered by the loss of Kv2.1, the size of individual RyR clusters was significantly reduced (Table 3). However, unlike the increased Cav1.2 immunolabeling found in adult Kv2.1 KO mouse brain sections, we found that neither the number nor size of somatic Cav1.2 clusters differed between WT and Kv2.1 KO CHNs. These observations suggests that while compensatory changes in Cav1.2 expression did not occur in cultured Kv2.1 KO CHNs after approximately two weeks *in vitro* as it did in adult brain neurons *in vivo*, the presence of Kv2.1 promoted the spatial coupling of Cav1.2 to RyRs.

**Table 3.**
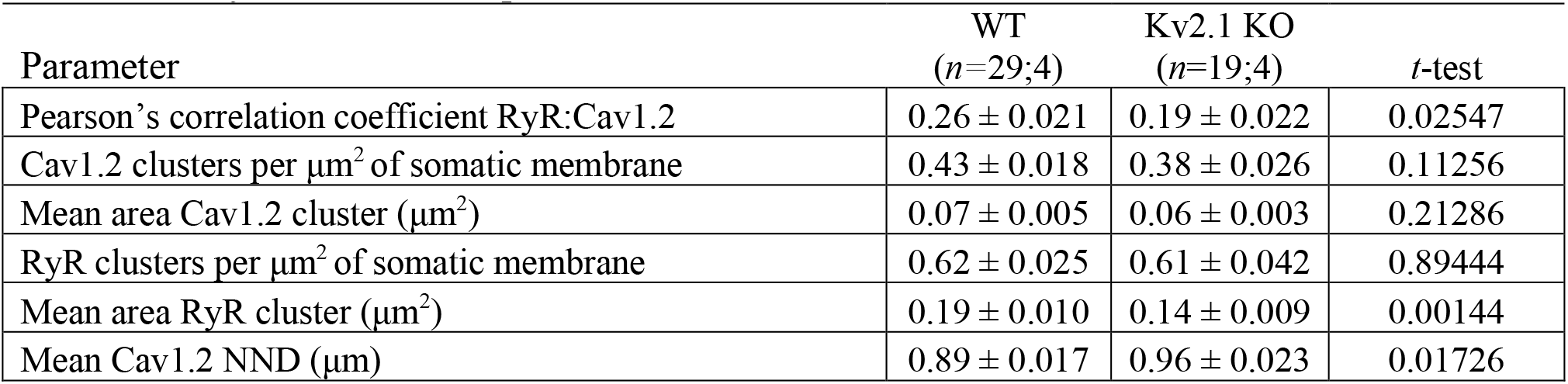
Cav1.2 and RyR colocalization parameters in mouse CH.

Finally, to evaluate how impaired Cav1.2 and RyR spatial coupling in Kv2.1 KO CHNs affected spontaneous CICR events or sparks, we imaged Cal-590-loaded cells using TIRF microscopy. Similar to rat CHNs, we observed spontaneous sparks in WT mouse CHNs that were associated with Kv2.1, Cav1.2, and RyR clusters identified by *post-hoc* immunolabeling (Fig. 8F). Consistent with the reduced colocalization of Cav1.2 and RyRs in Kv2.1 KO CHNs, we found that loss of Kv2.1 was associated with a significant reduction in spark frequency relative to WT control CHNs (Fig. 8G). Taken together, these findings demonstrate that Kv2.1 channels promote the spatial and functional association of endogenous Cav1.2 and RyRs in neurons, as well as the corresponding exogenous channels in HEK293T cells.

## Discussion

The findings in this study support a new model for the formation of Ca^2+^ signaling microdomains at EPJs and the local control of Ca^2+^ release from these structures. In this model, neuronal EPJs are Ca^2+^ signaling microdomains in which Cav1.2 and RyRs are brought into close proximity by Kv2.1-mediated clustering, forming a specialized somatic complex for the generation of localized Ca^2+^ signals by these Ca^2+^ channels. We propose that Kv2.1 channels function not only to anchor the ER to the PM but also to promote the organization of Cav1.2 channels into clusters in direct apposition to nearby ER RyRs. Our data indicate that Kv2.1-mediated clustering also increases the activity of Cav1.2. Spontaneous openings of Cav1.2 channels at negative potentials allow a small amount of Ca^2+^ to enter the cell at EPJs, activating nearby RyRs by the mechanism of CICR. The resulting Ca^2+^ sparks occur independently of action potentials. Thus, our model proposes a molecular structure underlying the localized somatodendritic Ca^2+^ signals previously observed in brain neurons (Berrout and Isokawa, 2009; Manita and Ross, 2009), and suggests a mechanism whereby Kv2.1 modulates these Ca^2+^ signals by simultaneously promoting the spatial association of Cav1.2 channels with RyRs and increasing their activity to trigger CICR.

### Kv2 channels dynamically cluster LTCCs

A key finding in this study is that endogenous LTCCs colocalized with clustered Kv2.1 in brain neurons, a finding supported by our crosslinking-based proteomic analyses showing that they exist in close spatial proximity. Moreover, colocalization of LTCCs with Kv2.1 could be reconstituted in heterologous cells, a property that required Kv2.1’s ability to cluster at EPJs but was separable from its voltage-gated K^+^ channel function. The Kv2.1-mediated association of Cav1.2 with EPJs appears to be dynamically regulated, as acute dispersal of Kv2.1 clusters in CHNs reduced Cav1.2’s association with RyRs and increased the nearest neighbor distance between individual Cav1.2 clusters. In addition, Kv2.1 expression in heterologous cells simultaneously enhanced the size of LTCC clusters and recruited LTCCs to Kv2.1-mediated EPJs. Consistent with this, we found that the spatial and functional coupling of somatic Cav1.2 channels to RyRs was reduced in Kv2.1 KO CHNs. Together, these findings indicate that LTCCs are recruited to Kv2.1-associated EPJs, a property we found was not shared by the T-type Ca^2+^ channel Cav3.1. Moreover, the co-purification of several Cavβ auxiliary subunits, which associate with LTCCs but not T-Type Ca^2+^ channels such as Cav3.1 (Fang and Colecraft, 2011), by IP of Kv2.1 from crosslinked brain samples further suggests a specific spatial interaction of LTCCs with Kv2.1. Numerous proteins have been identified that promote LTCC clustering in dendritic spines, including AKAP15 (Marshall et al., 2011) and PDZ domain-containing proteins (Zhang et al., 2005). The absence of these proteins from our proteomic analyses of proteins in close spatial proximity to Kv2.1, and our observation that expression of Kv2.1 increases clustering Cav1.2 in heterologous HEK293 cells, suggests that the proteins mediating Cav1.2 clustering in dendritic spines and at somatic EPJs may be distinct.

Although the molecular mechanism of Kv2.1 recruitment to EPJs is now established, and occurs via its phosphorylation-dependent interaction with VAPs (Johnson et al., 2018; Kirmiz et al., 2018b), the precise molecular mechanisms that underlies how LTCCs and RyRs are recruited to these sites is not yet clear. However, our data show that PM Cav1.2 organization was not impacted by coexpression of the clustering- and EPJ formation-deficient Kv2.1_S586A_ mutant as it was by Kv2.1_WT_ and the nonconducting Kv2.1_P404W_ point mutant. Additionally, Kv2.1_S586A_ was unable to enhance Cav1.2- and RyR-mediated sparks reconstituted in HEK293T cells, unlike these clustering-competent Kv2.1 isoforms. These findings suggest that Kv2.1 clustering and induction of EPJs is necessary for its spatial association with LTCCs.

The recruitment of LTCCs to EPJs in HEK293T cells formed upon heterologous expression of junctophilin-2 (Perni et al., 2017), an ER-localized protein critical for bridging the PM to the ER in myocytes (Jiang et al., 2016), is consistent with a model whereby tethering of LTCCs at or near Kv2-associated EPJs could be mediated by an intermediary recruited to Kv2.1-mediated EPJs, perhaps even one of the proteins identified in our proteomics analyses. We note that these proteomics analyses have the potential to identify proteins with lysine residues in close spatial proximity (≈12 Å) to those in Kv2.1, making them amenable to being crosslinked to Kv2.1 by DSP, and does not require their direct association. Moreover, the crosslinking reaction can yield “daisy-chained” protein linkages of spatially adjacent proteins. While at some point this would need to connect back to Kv2.1 to be immunopurified, the proteins present in the purified sample need not have this close spatial proximity to Kv2.1 itself.

However, it remains possible that PM Kv2s and LTCCs associate through a direct intermolecular interaction. Any domains on Kv2.1 contributing to this interaction would likely be conserved in Kv2.2, as we found that both channel paralogs similarly impacted LTCC cluster size and localization. It is unlikely that RyRs are directly recruited to EPJs by Kv2 channels, as RyR clusters persist in CHNs exposed to treatments that disperse Kv2.1 clusters (Misonou et al., 2005b) and while reduced in size in CA1 pyramidal neurons in the double Kv2.1/Kv2.2 knockout (Kirmiz et al., 2018a), in general RyR clusters persist in neurons in the brains of mice lacking Kv2 channels (Mandikian et al., 2014; Kirmiz et al., 2018a). Further experiments are needed to determine the molecular mechanisms and direct protein-protein interactions that result in the spatial association of these proteins at neuronal EPJs.

### Kv2.1-dependent potentiation of Cav1.2 currents

Given their prominent physiological role, the regulation of LTCCs is extensive and multimodal (Lipscombe et al., 2013; Hofmann et al., 2014; Neely and Hidalgo, 2014). The mechanisms involved in the modulation of LTCC function involve post-translational modification (*e.g.*, phosphorylation) or changes in the expression of the subunits (principal α1, and auxiliary Cavβ and α2δ) that together comprise the quaternary structure of an LTCC (Catterall, 2011; Zamponi et al., 2015). We have recently demonstrated a novel mechanism for regulating Cav1.2-(and Cav1.3-) containing LTCCs, whereby LTCCs function differently when clustered due to their clustering-dependent cooperative gating (Dixon et al., 2012; Dixon et al., 2015; Moreno et al., 2016). Thus, LTCC activity is sensitive to its spatial organization in the PM, influenced by its proximity to adjacent LTCCs (Navedo et al., 2005; Navedo et al., 2010; Dixon et al., 2012; Moreno et al., 2016) and also to its localization to specific neuronal compartments (Hall et al., 2013; Tseng et al., 2017). In neurons, such regulation likely acts to ensure that Cav1.2 is most active when properly targeted to specific subcellular domains and less active when outside these regions. Here, we show that the subcellular localization and activity of somatic Cav1.2 channels are influenced by Kv2.1, which increases both Cav1.2 clustering and its opening at polarized *V*_m_ values. At least two other proteins, α-actinin (Hall et al., 2013) and densin-180 (Wang et al., 2017), exert a similar dual regulation on neuronal Cav1.2, by promoting its localization to dendritic spines and enhancing its activity at these sites. Neither of these proteins was identified in our proteomic analyses of proteins in close spatial proximity to Kv2.1, further suggesting that Cav1.2 complexes in dendritic spines and at somatic EPJs may be distinct. The reduced whole-cell LTCC currents and impaired association of somatic Cav1.2 with RyRs in Kv2.1 KO CHNs suggests that Kv2.1 serves this dual targeting/modulation function for LTCCs within the soma and proximal dendrites.

In both CHNs and HEK293T cells, currents resulting from the opening of endogenous and exogenous Cav1.2 channels, respectively, are increased in the presence of Kv2.1. In HEK293T cells, Cav1.2 channels coexpressed with clustered Kv2.1 are activated at more polarized *V*_m_ values relative to those produced by Cav1.2 alone. The Kv2.1-dependent increase in whole-cell Cav1.2 current amplitude in both cell types occurs without an apparent change in the number of Cav1.2 channels present on the PM, as total Cav1.2 on-gating charges were unaltered by coexpression with Kv2.1. Instead, it appears that coupling of Cav1.2 voltage sensor movement to channel opening is enhanced in the presence of Kv2.1. What is the molecular mechanism underlying this effect on Cav1.2 channel opening? We suggest three possibilities. First, the increase in *I*_Ca_ and leftward shift in the voltage-dependence of activation that we observed upon coexpression of Kv2.1 in HEK293T cells are similar to those observed during optogenetic induction of Cav1.2 channel oligomerization. Thus, one possible mechanism is that Kv2.1-induced clustering at EPJs increases the probability of physical interactions between Cav1.2 channels, which promotes their cooperative gating (Dixon et al., 2012; Navedo et al., 2010). A second possibility is that Kv2.1 functions as an auxiliary voltage sensor for Cav1.2 channels, perhaps through a direct intermolecular interaction of the two channels. However, the apparent localization of many Cav1.2 clusters adjacent to rather than directly overlapping with Kv2.1 clusters in CHNs (e.g., see Fig. 1D-E) suggests that although these proteins associate in close spatial proximity, there may not be a direct interaction between individual Kv2.1 and Cav1.2 channels.

A third potential explanation for the Kv2.1-mediated increase in Cav1.2 channel activity is that Cav1.2 is modulated by signaling molecules recruited to EPJs by Kv2.1. It is well established that phosphorylation of Cav1.2 is a major mechanism to regulate its activity. Phosphorylation by protein kinase A (PKA) increases Ca^2+^ influx through Cav1.2, enhancing CICR (Dittmer et al., 2019). Another candidate which might impact Cav1.2 at EPJs is Ca^2+/^calmodulin-dependent protein kinase II (CaMKII), which has also been shown to interact with Kv2.1 (McCord and Aizenman, 2013). Enhanced Cav1.2 opening at polarized *V*_m_ values and increased open probability are produced by both PKA-(Tsien et al., 1986; Bers and Perez-Reyes, 1999) and CaMKII-(Erxleben et al., 2006; Blaich et al., 2010) dependent phosphorylation of Cav1.2. Moreover, given the well-established association of RyRs with PKA and CaMKII (Zalk et al., 2007), it is conceivable that RyRs, Cav1.2, and Kv2.1 are substrates of these protein kinases at somatic EPJs. A recent study showed that in dendritic EPJs adjacent to spines, Cav1.2 is inhibited through a direct interaction with the ER-localized protein stromal interaction molecule 1 (STIM1) in a negative feedback response to Cav1.2- and RyR-mediated CICR (Dittmer et al., 2019). As such the Kv2.1-mediated localization of Cav1.2 at EPJs may bring it in close proximity to numerous regulatory molecules, at least a subset of which may also be expressed in HEK293T cells in which we also observed prominent effects of Kv2.1 clustering on Cav1.2 activity.

### Properties of Ca^2+^ sparks at Kv2.1-associated EPJs

The results presented here indicate that Ca^2+^ sparks occurring at Kv2.1-associated EPJs were triggered primarily by Ca^2+^ influx through LTCCs initiating the opening of juxtaposed RyRs. Accordingly, Ca^2+^ spark frequency increased when neurons were exposed to Cav1.2 channel agonists and decreased by blockade of LTCCs. Indeed, loss of Kv2.1 expression was associated with a decrease in Ca^2+^ spark frequency, likely because of decreased spatial association of Cav1.2 and RyRs, decreased RyR cluster size, and decreased LTCC currents.

Our findings indicate that Kv2.1-mediated somatodendritic EPJs provide a molecular platform to elevate local Ca^2+^ at individual EPJs without an increase in global Ca^2+^, but that can also contribute to global, action potential-induced increases in cytoplasmic Ca^2+^. These results reinforce previous observations (Berrout and Isokawa, 2009; Manita and Ross, 2009; Miyazaki et al., 2012; Miyazaki and Ross, 2013) that hippocampal neurons possess the molecular machinery to produce spontaneous local elevations in somatodendritic Ca^2+^ that could potentially impact a wide variety of signaling pathways.

That sparks can occur independently in neighboring Kv2.1-containing EPJs suggests a mechanism for compartmentalized Ca^2+^ signaling in the aspiny regions of neurons (somata, proximal dendrites, axon initial segment) in which Kv2.1 clusters are located. One specific role identified for Ca^2+^ signals produced by somatic RyR receptors at EPJs is in cartwheel cells (inhibitory interneurons found in the dorsal cochlear nucleus), where they trigger rapid gating of BK Ca^2+^-activated K^+^ channels to control electrical excitability (Irie and Trussell, 2017). While this mode of BK channel activation has not been observed in CA1 pyramidal neurons (Ross, 2012), sparks at Kv2.1-associated EPJs might influence electrical activity in pyramidal cells through Ca^2+^-sensitive enzymes that modify ion channel function, such as protein kinases and phosphatases that influence their phosphorylation state (Misonou et al., 2004). In addition, a role for somatic Ca^2+^ sparks has been identified in DRG neurons, where they promote non-synaptic exocytosis of ATP-loaded secretory vesicles (Ouyang et al., 2005). Whether Ca^2+^ entry mediated by LTCCs and RyRs at Kv2.1-associated EPJs impacts secretory vesicle exocytosis in brain neurons will need to be investigated in future studies.

### Potential impact on downstream signaling pathways

Somatodendritic LTCCs are preferentially coupled to activation of signaling pathways resulting in changes in gene expression (Wheeler et al., 2012; Wild et al., 2019). In sympathetic neurons, local Ca^2+^ influx through LTCCs rather than bulk elevation of intracellular Ca^2+^ efficiently activates the transcription factor cAMP response element–binding protein (CREB) (Wheeler et al., 2008) through a mechanism that involves a signaling complex containing components of a PM-to-nucleus Ca^2+^ shuttle (Ma et al., 2012; Ma et al., 2014; Cohen et al., 2015). Moreover, somatic LTCCs play a unique role in the Ca^2+^ influx that leads to activation of the NFAT transcription factor (Wild et al., 2019). The results presented here suggest that Kv2.1-mediated organization and regulation of somatic LTCCs provides a molecular mechanism to control local Ca^2+^ influx and serve as an organizer of Ca^2+^ signaling microdomains. Previous work from us (Misonou et al., 2004; Misonou et al., 2005a) and others (Mulholland et al., 2008; Aras et al., 2009) has shown that acute ischemic or depolarizing events disperse Kv2.1 clusters and polarize its *V*_m_ activation threshold, potentially as a homeostatic mechanism to reduce neuronal activity and Ca^2+^ overload that can lead to excitotoxicity. In our experiments here, we determined that Kv2.1-mediated clustering was associated with enhanced functional coupling of Cav1.2 and RyRs, as well as increased activation of Cav1.2 at polarized *V*_m_ values. Therefore, dispersal of Kv2.1 clusters and the resulting dissociation of Cav1.2 and RyRs may represent a negative feedback loop to limit excessive increases in cytoplasmic Ca^2+^. By decreasing LTCC- and RyR-mediated CICR, dispersal of Kv2.1 clusters may help to curb excessive accumulation of intracellular Ca^2+^, which inappropriately activates signaling pathways contributing to neuronal damage or death (Dirnagl et al., 1999). Activity-dependent declustering of Kv2.1 may also help to reduce currents conducted by LTCCs, both through increased activation of hyperpolarizing Kv2.1 currents at polarized *V*_m_ (opposing activation of voltage-gated Ca^2+^ channels) and also through limiting Cav1.2 activity by altering its spatial organization in the PM. Our findings may also contribute to an understanding of the pathogenic mechanisms underlying mutations in Kv2.1 predicted to selectively disrupt the PRC domain required for Kv2.1 clustering (de Kovel et al., 2017).

Overall, the findings presented here identify a molecular structure underlying the spontaneous somatodendritic Ca^2+^ signals previously observed in hippocampal pyramidal neurons. While our live cell experiments were primarily confined to CHNs cultured for 9-15 days *in vitro*, our data indicate that the spatial association of Kv2.1, Cav1.2, and RyRs is preserved in intact adult mouse and rat brains and can be recapitulated in heterologous cells. Moreover, somatodendritic Ca^2+^ sparks have been observed in acute hippocampal slices obtained from rats aged P3-P80 (Miyazaki et al., 2012), suggesting that these Ca^2+^ release events serve functional roles that emerge early in pyramidal neuron development and continue beyond this period. Although it is unclear whether spontaneous Ca^2+^ sparks serve a specific function at their site of generation, or if they instead reflect stochastic events whose primary impact lies in their group behavior (i.e., through modulation of bulk cytosolic Ca^2+^), the results described here have relevance to obtaining a better understanding of their generation as well as their downstream effects.

## Materials and Methods

### Animals

All procedures involving rats and mice were approved by the University of California, Davis Institutional Animal Care and Use Committee and performed in accordance with the NIH Guide for the Care and Use of Laboratory Animals. Animals were maintained under standard light-dark cycles and allowed to feed and drink ad libitum. Sprague-Dawley rats were used for immunolabeling experiments and as a source of hippocampal neurons for primary culture. Kv2.1 KO mice (RRID:IMSR_MGI:3806050) (Jacobson et al., 2007; Speca et al., 2014) were generated from breeding of *Kcnb1*^+/-^ mice that had been backcrossed on the C57/BL6J background (RRID:IMSR_JAX:000664). Littermates were used when available. Adult male (mice and rats) and female (rats) were used in immunohistochemistry experiments; adult male and female mice were used in proteomics. Experiments using CHNs were performed using neuronal cultures obtained from pooling neurons from animals of both sexes (rats and mice) and also cultures in which individual pups were grouped by sex after visual inspection (mice).

### Antibodies

**Table 4.**
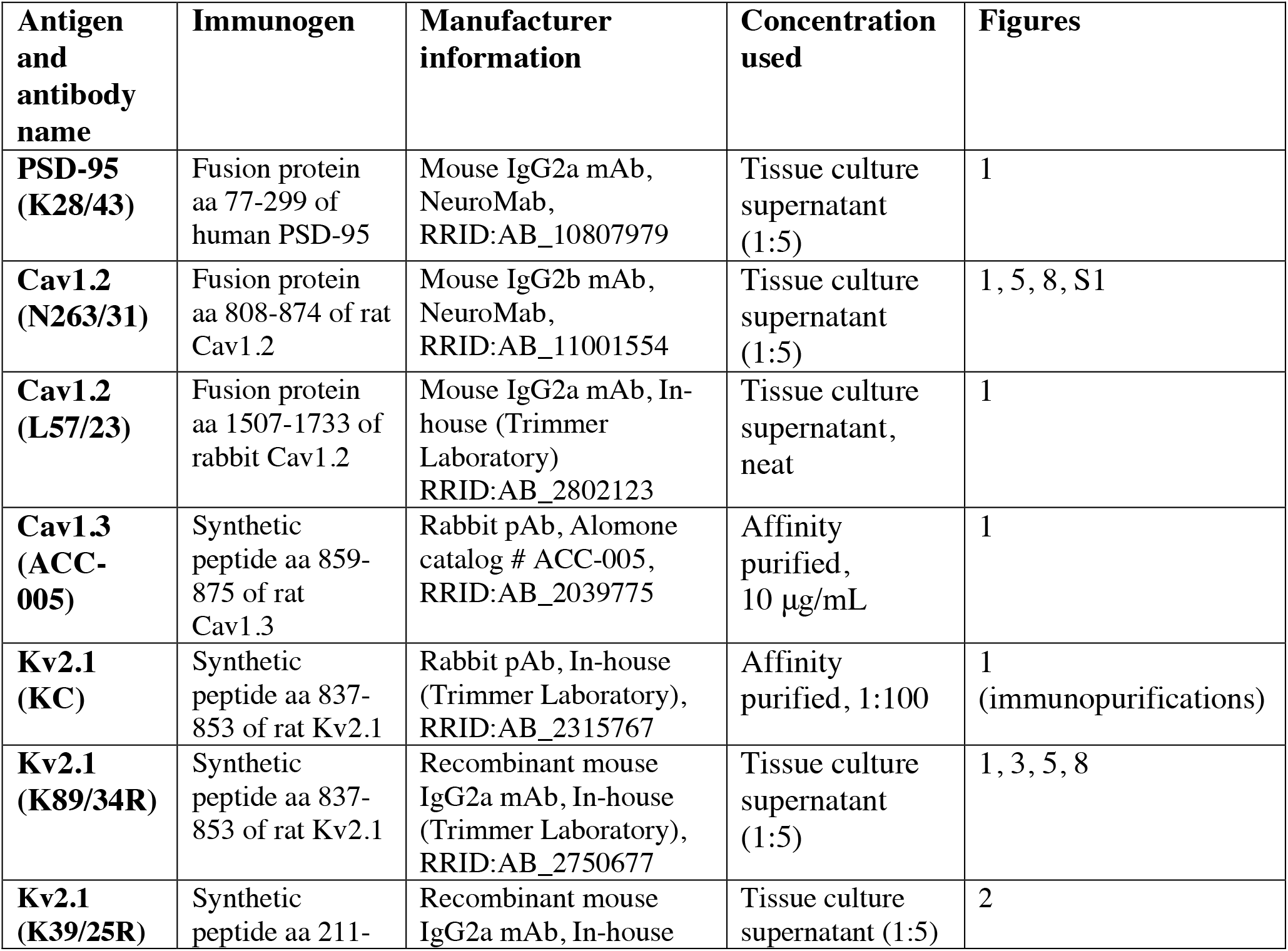
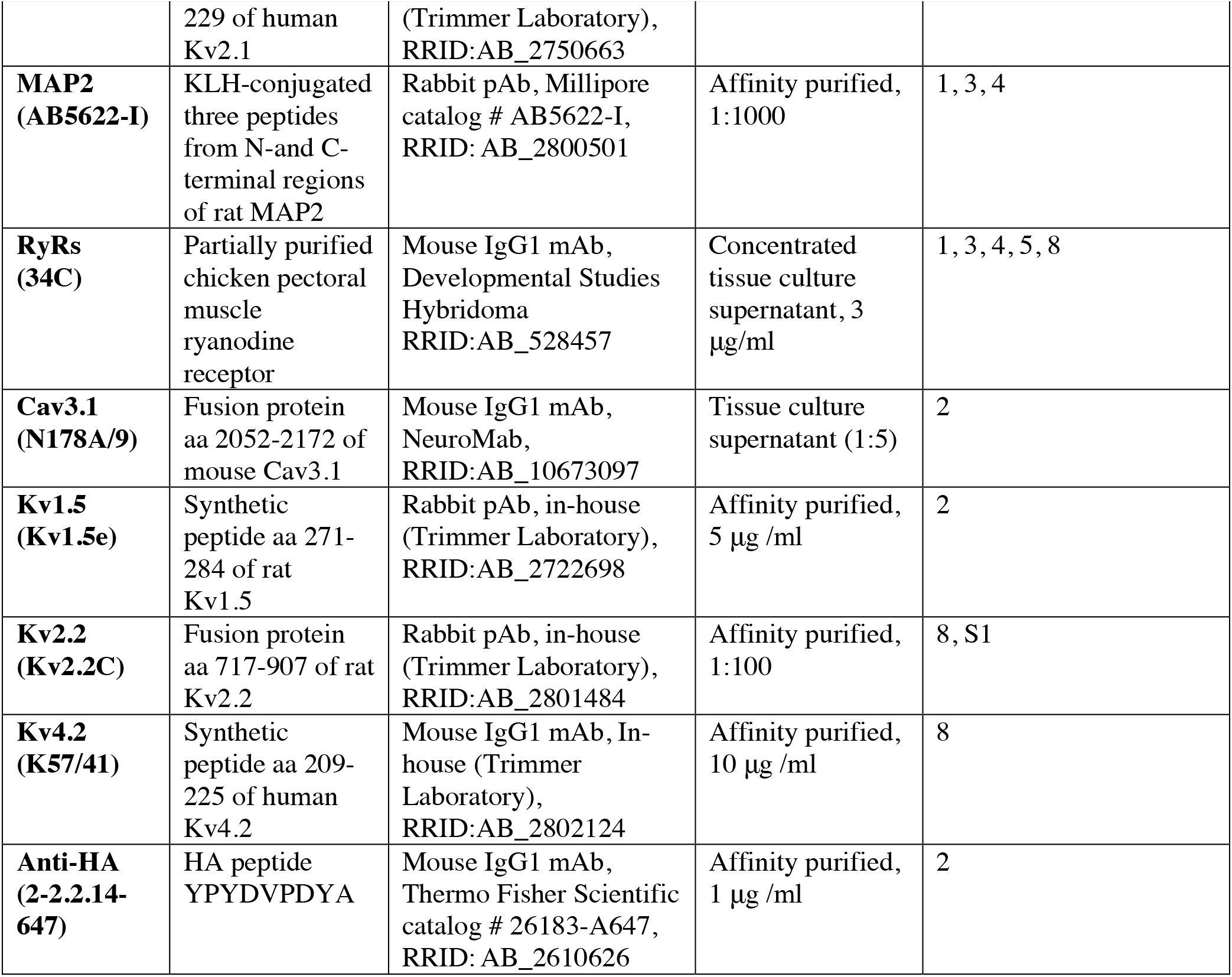
Antibody information.

### Hippocampal neuron cultures

Neuronal cultures were prepared and maintained as previously described (Kirmiz et al., 2018a; Kirmiz et al., 2018b). Hippocampi were dissected from either postnatal day 0-1 pups (mice) following genotyping or embryonic day 18 embryos (rat) and dissociated enzymatically for 20 min at 37 °C in HBSS supplemented with 0.25% (w/v) trypsin (Worthington Cat# LS003707), followed by mechanical dissociation via trituration with fire-polished glass Pasteur pipettes. Dissociated cells were suspended in plating medium containing Neurobasal (ThermoFisher Cat# 21103049) supplemented with 10% fetal bovine serum (FBS, Invitrogen Cat# 16140071), 2% B27 supplement (Invitrogen Cat# 17504044), 2% GlutaMAX (ThermoFisher Cat# 35050061), and 0.001% gentamycin (Gibco Cat# 15710064) and plated at 60,000 cells per dish in glass bottom dishes (MatTek Cat# P35G-1.5-14-C) or on microscope cover glasses (Karl Hecht Assistent Ref# 92099005050) coated with poly-L-lysine (Sigma Cat# P2636). After 5 days in vitro (DIV), cytosine-D-arabinofuranoside (Millipore Cat# 251010) was added to inhibit non-neuronal cell growth. Neurons were transiently transfected at DIV 7-10 using Lipofectamine 2000 (Invitrogen Cat# 11668019) for 1.5 hours as previously described (Lim et al., 2000). Neurons were used for experiments 40-48 hours post transfection.

For glutamate-induced dispersal of Kv2.1 in rat CHNs, 20-24 DIV neurons cultured on microscope cover glasses were incubated in 1 mL of a modified Krebs-Ringer buffer (KRB) containing (in mM): 146 NaCl, 4.7 KCl, 2.5 CaCl_2_, 0.6 MgSO_4_, 1.6 NaHCO_3_, 0.15 NaH_2_PO_4_, 8 glucose, 20 HEPES, pH 7.4, approximately 330 mOsm for 30 minutes at 37 °C. We then added an additional 1 mL of KRB prewarmed to 37 °C, with or without 20 µM glutamate (Calbiochem Cat #3510) for a final glutamate concentration of 10 µM, and incubated CHNs for 10 minutes at 37 °C. We then proceeded immediately to fixation.

### HEK293T cell culture

HEK293T cells (ATCC Cat# CRL-3216) were maintained in Dulbecco’s modified Eagle’s medium (Gibco Cat# 11995065) supplemented with 10% Fetal Clone III (HyClone Cat# SH30109.03), 1% penicillin/streptomycin, and 1x GlutaMAX (ThermoFisher Cat# 35050061) in a humidified incubator at 37 °C and 5% CO_2_. Cells were transiently transfected using Lipofectamine 2000 following the manufacturer’s protocol, in DMEM without supplements, then returned to regular growth medium 4 hours after transfection. 20-24 hours later, cells were passaged to obtain single cells on glass bottom dishes (MatTek Cat# P35G-1.5-14-C) or microscope cover glasses (VWR Cat# 48366-227) coated with poly-L lysine. Cells were then used for experiments approximately 15 hours after being passaged.

### Immunolabeling of cells

Immunolabeling of CHNs and HEK293T cells was performed as described previously (Kirmiz et al., 2018b). CHNs were fixed in ice cold 4% (wt/vol) formaldehyde (freshly prepared from paraformaldehyde, Fisher Cat# O4042) in phosphate buffered saline (PBS, Sigma Cat #P3813), pH 7.4, for 15 minutes at 4 °C. HEK293T cells were fixed in 3.2% formaldehyde (freshly prepared from paraformaldehyde) and 0.1% glutaraldehyde (Ted Pella, Inc., Cat# 18426) prepared in PBS pH 7.4, for 20 minutes at room temperature (RT), washed 3 x 5 min in PBS and quenched with 1% sodium borohydride (Sigma Cat# 452882) in PBS for 15 minutes at room temperature. All subsequent steps were performed at RT. Cells were then washed 3 x 5 minutes in PBS, followed by blocking in blotto-T (Tris-buffered saline [10 mM Tris, 150 mM NaCl, pH 7.4] supplemented with 4% (w/v) non-fat milk powder and 0.1 % (v/v) Triton-X100 [Roche Cat# 10789704001]) for 1 hour. Cells were immunolabeled for 1 hour with primary antibodies diluted in blotto-T (concentrations used for primary antibodies listed in Table 4). Following 3 x 5 minute washes in blotto-T, cells were incubated with mouse IgG subclass- and/or species-specific Alexa-conjugated fluorescent secondary antibodies (Invitrogen) diluted in blotto-T for 45 min, then washed 3 x 5 minutes in PBS. Cover glasses were mounted on microscope slides with Prolong Gold mounting medium (ThermoFisher Cat # P36930) according to the manufacturer’s instructions. For surface immunolabeling of HEK293T cells, cells were fixed for 20 minutes at 4 °C in ice-cold 3.2% formaldehyde prepared in PBS, pH 7.4. All subsequent steps were performed at RT without Triton X-100. Cells were washed 3 x 10 minutes in PBS, blocked for 1 h in blotto-T, then incubated for 2 hours in primary antibodies diluted in blotto-T without Triton X-100. Cells were then washed 3 x 15 minutes in blotto-T, then incubated with mouse IgG subclass- and/or species-specific Alexa-conjugated fluorescent secondary antibodies diluted in blotto-T for 1.5 hours. Cells were washed 3 x 15 minutes in PBS, then mounted onto microscope slides as described above.

Widefield fluorescence images were acquired with an AxioCam MRm digital camera installed on a Zeiss AxioImager M2 microscope or with an AxioCam HRm digital camera installed on a Zeiss AxioObserver Z1 microscope with a 63X/1.40 NA plan-Apochromat oil immersion objective and an ApoTome coupled to Axiovision software (Zeiss, Oberkochen, Germany). Confocal images were acquired using a Zeiss LSM880 confocal laser scanning microscope equipped with an Airyscan detection unit and a Plan-Apochromat 63x/1.40 NA Oil DIC M27 objective. Structured illumination microscopy (N-SIM) images were acquired with a Hamamatsu ORCA-ERCCD camera on a SIM/wide-field capable Nikon Eclipse Ti microscope with an EXFO X-Cite metal halide light source and a 100X PlanApo TIRF/1.49 objective. Colocalization and morphological analyses of Cav1.2 and RyRs in CHNs was performed using FIJI (NIH). For the colocalization analyses, an ROI was drawn around the soma of a neuron and PCC values were collected using the Coloc2 plugin. All intensity measurements were collected using FIJI. All intensity measurements reported in line scans are normalized to the maximum intensity measurement. Measurements of cluster sizes was performed essentially as previously described (Kirmiz et al., 2018a; Kirmiz et al., 2018b). Images were subjected to rolling ball background subtraction and subsequently converted into a binary mask by thresholding. Cluster sizes were measured using the “analyze particles” feature of FIJI; nearest neighbor distances were calculated from cluster centroid values using the nearest neighbor distance plugin in FIJI. For presentation, images were exported as TIFFs and linearly scaled for min/max intensity and flattened as RGB TIFFs in Photoshop (Adobe).

### Immunolabeling of brain sections

Following administration of pentobarbital to induce deep anesthesia, animals were transcardially perfused with 4% formaldehyde (freshly prepared from paraformaldehyde) in 0.1 M sodium phosphate buffer pH 7.4 (0.1 M PB). Sagittal brain sections (30 µm thick) were prepared and immunolabeled using free-floating methods as detailed previously (Rhodes et al., 2004; Speca et al., 2014; Bishop et al., 2015; Palacio et al., 2017). Sections were permeabilized and blocked in 0.1 M PB containing 10% goat serum and 0.3% Triton X-100 (vehicle) for 1 hour at RT, then incubated overnight at 4°C in primary antibodies (Table 4) diluted in vehicle. After 4 x 5 minute washes in 0.1 M PB, sections were incubated with mouse IgG subclass- and/or species-specific Alexa-conjugated fluorescent secondary antibodies (Invitrogen) and Hoechst 33258 DNA stain diluted in vehicle at RT for 1 hour. After 2 x 5 minute washes in 0.1 M PB followed by a single 5 minute wash in 0.05 M PB, sections were mounted and air dried onto gelatin-coated microscope slides, treated with 0.05% Sudan Black (EM Sciences) in 70% ethanol for 2 minutes (Schnell et al., 1999). Samples were then washed extensively in water and mounted with Prolong Gold (ThermoFisher Cat # P36930). Images of brain sections were taken using the same exposure time to compare the signal intensity directly using an AxioCam HRm high-resolution CCD camera installed on an AxioObserver Z1 microscope with a 10x/0.5 NA lens, and an ApoTome coupled to Axiovision software, version 4.8.2.0 (Zeiss, Oberkochen, Germany). Labeling intensity within stratum pyramidale and stratum radiatum of hippocampal area CA1 was measured using a rectangular region of interest (ROI) of approximately 35 μm x 185 μm. Labeling intensity within stratum granulosum and the inner third of stratum moleculare of the dentate gyrus (DG) was measured using a rectangular ROI of approximately 48 μm x 200 μm. To maintain consistency between samples, the average pixel intensity values of ROIs from CA1 were acquired near the border of CA1 and CA2, and those from DG were obtained near the center of the dorsal/suprapyramidal blade of the DG. Signal intensity values from all immunolabels and of Hoechst dye were measured from the same ROI. Background levels for individual labels were measured from no primary controls for each animal and subtracted from ROI values. High magnification confocal images of rat and mouse hippocampus were acquired using a Zeiss LSM880 confocal laser scanning microscope equipped with an Airyscan detection unit and a Plan-Apochromat 63x/1.40 NA Oil DIC M27 objective.

### Immunopurification of Kv2.1 and proteomics

Crosslinked mouse brain samples for immunopurification were prepared as previously described (Kirmiz et al., 2018b). Individual excised brains from three wild-type and three Kv2.1 KO mouse littermates were homogenized separately over ice in a Dounce homogenizer containing 5 mL homogenization and crosslinking buffer (in mM): 320 sucrose, 5 NaPO_4_, pH 7.4, supplemented with 100 NaF, 1 PMSF, protease inhibitors, and 1 DSP (Lomant’s reagent, ThermoFisher Cat# 22585). Following a 1 hour incubation on ice, DSP was quenched with 20 mM Tris, pH 7.4 (JT Baker Cat# 4109-01 [Tris base]; and 4103-01 [Tris-HCl]). 2 mL of each homogenate was then added to an equal volume of ice-cold 2x radioimmunoprecipitation assay (RIPA) buffer (final concentrations): 1% (vol/vol) TX-100, 0.5% (wt/vol) deoxycholate, 0.1% (wt/vol) SDS, 150 NaCl, 50 Tris, pH 8.0 and incubated on a tube rotator at 4 °C for 30 minutes. Insoluble material was then pelleted by centrifugation at 12,000 × g for 10 minutes at 4 °C. The supernatants from the six brains were incubated overnight at 4 °C with the anti-Kv2.1 rabbit polyclonal antibody KC (Trimmer, 1991). Following this incubation, we added 100 μL of magnetic protein G beads (ThermoFisher Cat# 10004D) and incubated the samples on a tube rotator at 4 °C for 1 h. Beads were then washed 6x following capture on a magnet in ice-cold 1x RIPA buffer, followed by four washes in 50 mM ammonium bicarbonate (pH 7.4). Proteins captured on magnetic beads were digested with 1.5 mg/mL trypsin (Promega Cat# V5111) in 50 mM ammonium bicarbonate overnight at 37 °C. The eluate was then lyophilized and resuspended in 0.1% trifluoroacetic acid in 60% acetonitrile.

Proteomic profiling was performed at the University of California, Davis Proteomics Facility. Tryptic peptide fragments were analyzed by LC-MS/MS on a Thermo Scientific Q Exactive Plus Orbitrap Mass spectrometer in conjunction with a Proxeon Easy-nLC II HPLC (Thermo Scientific) and Proxeon nanospray source. Digested peptides were loaded onto a 100 μm x 25 mm Magic C18 100Å 5U reverse phase trap where they were desalted online, then separated using a 75 μm x 150 mm Magic C18 200Å 3U reverse phase column. Peptides were eluted using a 60-minute gradient at a flow rate of 300 nL per min. An MS survey scan was obtained for the *m/z* range 350-1600; tandem MS spectra were acquired using a top 15 method, where the top 15 ions in the MS spectrum were subjected to HCD (High Energy Collisional Dissociation). Precursor ion selection was performed using a mass window of 1.6 *m/z*, and normalized collision energy of 27% was used for fragmentation. A 15 s duration was used for the dynamic exclusion. MS/MS spectra were extracted and charge state deconvoluted by Proteome Discoverer (Thermo Scientific). MS/MS samples were then analyzed using X! Tandem (The GPM, thegpm.org; version Alanine (2017. 2. 1.4)). X! Tandem compared acquired spectra against the UniProt Mouse database (May 2017, 103089 entries), the cRAP database of common proteomic contaminants (www.thegpm.org/crap; 114 entries), the ADAR2 catalytic domain sequence, plus an equal number of reverse protein sequences assuming the digestion enzyme trypsin. X! Tandem was searched with a fragment ion mass tolerance of 20 ppm and a parent ion tolerance of 20 ppm. Variable modifications specified in X! Tandem included deamidation of asparagine and glutamine, oxidation of methionine and tryptophan, sulfone of methionine, tryptophan oxidation to formylkynurenin of tryptophan and acetylation of the N-terminus. Scaffold (version Scaffold_4.8.4, Proteome Software Inc., Portland, OR) was used to validate tandem MS-based peptide and protein identifications. X! Tandem identifications were accepted if they possessed -Log (Expect Scores) scores of greater than 2.0 with a mass accuracy of 5 ppm. Protein identifications were accepted if they contained at least two identified peptides. The threshold for peptide acceptance was greater than 95% probability. After filtering for proteins present in the wild-type brain samples and absent in the KO brain samples, proteins in the wild-type sample were ranked by spectral counts. To generate the data in Table 2, spectral counts for the individual proteins in the three separate IPs were then normalized to the spectral counts for Kv2.1 (set at 100).

### Plasmid constructs

To maintain consistency with previous studies, we use the original (Frech et al., 1989) amino acid numbering of rat Kv2.1 (accession number NP_037318.1). The generation of DsRed-Kv2.1 and -Kv2.2 plasmids has been described previously (Kirmiz et al., 2018b). GCaMP3-Kv2.1 was generated using Gibson assembly to insert GCaMP3 (Tian et al., 2009) into the Kv2.1 RBG4 vector (Shi et al., 1994), resulting in fusion of GCaMP3 to the N-terminus of full-length rat Kv2.1. The plasmid encoding Kv2.1_S586A_ has been previously described (Lim et al., 2000); the plasmid encoding Kv2.1_P404W_ in the pcDNA4/TO vector was a gift from Dr. Jon Sack (University of California, Davis). The plasmid encoding Kv1.5 has been previously described (Nakahira et al., 1996). The plasmids encoding GFP-tagged full-length rabbit Cav1.2 α1 subunit (accession number NP_001129994.1) and GFP-tagged short isoform of rat Cav1.3 α subunit (accession AAK72959.1) have been previously described (Dixon et al., 2015; Moreno et al., 2016). Plasmids encoding untagged full-length mouse Cav1.2, rat Cavβ3, and rat α_2_δ_1_ were gifts of Dr. Diane Lipscombe (Brown University). The plasmid encoding BFP-Sec61β was a gift from Dr. Jodi Nunnari (University of California, Davis). Plasmid encoding HA-tagged rat Cav1.2 was a gift from Dr. Valentina Di Biase (Medical University of Graz), plasmid encoding human Cav3.1 was a gift from Dr. Edward Perez-Reyes (University of Virginia), and plasmid encoding full-length mouse RyR2 fused with YFP was a gift of Dr. Alla Fomina (University of California, Davis). The vector encoding human STAC1 was obtained from DNASU (DNASU plasmid # HsCD00445396).

### Live cell imaging

HEK293T cells transfected with RyR2-YFP, LTCC α1 subunit (Cav1.2 or Cav1.3(s)), Cavβ3, Cavα_2_δ_1_, STAC1, and empty vector control (pcDNA3) or DsRed-Kv2.1_P404W_ plasmids in a 1.5:1:0.5:0.5:0.25:1 ratio were seeded to glass bottom dishes (MatTek Cat# P35G-1.5-14-C) approximately 15 hours prior to recording. Total internal reflection fluorescence (TIRF) and widefield microscopy imaging of HEK293T cells and DIV9-10 (transfected with GCaMP3-Kv2.1) or DIV14-21 (loaded with Cal-590 AM) CHNs cultured on glass-bottom dishes was performed in KRB at 37 °C as previously described (Kirmiz et al., 2018b). For imaging of cells loaded with Ca^2+^-sensitive dye, cells were first incubated in regular culture medium to which had been added 1.5 µM Cal-590 AM (AAT Bioquest Cat# 20510) for 45 minutes or Fluo-4 AM (Invitrogen Cat# F14201) for 25 minutes at 37 °C. Dye-containing medium was then aspirated, followed by two washes in KRB which had been warmed to 37 °C. Cells were then incubated in KRB for an additional 30 minutes at 37 °C prior to imaging. Caffeine (Sigma Cat# C0750), thapsigargin (Millipore Cat# 586005), nimodipine (Alomone Cat# N-150), Bay K8644 (Alomone Cat# B-350), and tetracaine (Sigma Cat# T7508) were dissolved in warm KRB at 2x the final concentration and added to cells during imaging by a pipette. Images were acquired on a Nikon Eclipse Ti TIRF/widefield microscope equipped with an Andor iXon EMCCD camera and a Nikon LUA4 laser launch with 405, 488, 561, and 647 nm lasers, using a 100x/1.49 NA PlanApo TIRF objective and NIS Elements software. For *post-hoc* immunolabeling of CHNs, the dish orientation and location of the imaged cell was recorded, after which the CHN was fixed in ice-cold 4% formaldehyde and processed for immunolabeling as described above. Recorded CHNs were identified on the basis of expression of GCaMP3-Kv2.1 and/or neurite morphology revealed by immunolabeling for MAP2.

### Electrophysiology

HEK293T cells transfected with Cav1.2-GFP, Cavβ3, Cavα_2_δ_1_, and empty vector control (pcDNA3) or DsRed-Kv2.1_P404W_ plasmids in a 1:0.5:0.5:1 ratio were seeded to microscope cover glasses (Fisher Cat# 12-545-102) approximately 15 hours prior to recording to obtain single cells. Coexpression of Cav1.2 and Kv2.1_P404W_ in HEK293T cells was apparently cytotoxic and thus necessitated seeding of cells at a higher density to obtain viable single cells as compared to control cells expressing Cav1.2 alone. HEK293T cells were patched in an external solution of modified Krebs-Ringer buffer (KRB) containing (in mM): 146 NaCl, 4.7 KCl, 2.5 CaCl_2_, 0.6 MgSO_4_, 1.6 NaHCO_3_, 0.15 NaH_2_PO_4_, 8 glucose, 20 HEPES, pH 7.4, approximately 330 mOsm. Transfected cells were identified by the presence of GFP and DsRed expression. *I*_Ca_ was recorded in transfected cells using the whole-cell voltage clamp patch configuration using fire-polished borosilicate pipettes that had resistances of 2-3 MΩ when filled with an internal solution containing (in mM): 125 Cs-methanesulfonate, 10 TEA-Cl, 1 MgCl_2_, 0.3 Na_2_-GTP, 13 phosphocreatine-(di)Tris, 5 Mg·ATP, 5 EGTA, 10 HEPES, adjusted to pH 7.22 with CsOH, approximately 320 mOsm. Currents were sampled at 20 kHz and low-pass–filtered at 2 kHz using an Axopatch 200B amplifier, and acquired using pClamp 10.2 software (Molecular Devices, Sunnyvale, CA). All experiments were performed at room temperature (22–25 °C). Pipette capacitance was compensated using the amplifier, and capacitance and ohmic leak were subtracted online using a P/5 protocol. Current–voltage (*I*–*V*) relationships were obtained approximately three minutes after obtaining the whole-cell configuration by subjecting cells to a series of 300-ms depolarizing pulses from the holding potential of -70 mV to test potentials ranging from -60 to +100 mV in 10 mV increments. The voltage dependence of *G*/*G*_max_ was obtained from the recorded currents by converting them to conductances (*G*) using the equation *G* = *I*_Ca_/(test pulse potential – *E*_rev(Ca)_), plotting the normalized values (*G*/*G*_max_) versus the test potential, and fitting them to a Boltzmann function. Steady-state inactivation was measured by subjecting cells to a series of 2500-ms conditioning prepulses from the holding potential to potentials ranging from -60 to +100 mV, returning to the -70 mV holding potential for 5 ms, then measuring the peak current elicited by a 300 ms step to the -20 mV test potential. Data was analyzed and plotted using Prism software (Graphpad Software Inc., San Diego, CA). For experiments in which depolarization-induced increases in Ca^2+^-sensitive dye were measured, we included 0.2 mM Rhod-2 (AAT Bioquest Cat# 21068) in the patch pipette solution. Images were acquired at 10 Hz using a through-the-lens TIRF microscope built around an Olympus IX-70 inverted microscope equipped with an oil-immersion ApoN 60x/1.49 NA TIRF objective and an Andor iXON CCD camera using TILLvisION imaging software (TILL Photonics, FEI, Hillsboro, OR).

To measure gating and ionic tail currents, we first determined the reversal potential for *I*_Ca_ from the *I–V* relationship obtained using the *I–V* protocol described above. Gating currents were then measured by applying a series of depolarizing steps from the holding potential (-70 mV) to potentials ±5 mV of the reversal potential in 1 mV increments. Currents were sampled at a frequency of 25 kHz and low-pass filtered at 2 kHz. We first obtained recordings in cells perfused with KRB alone, then obtained recordings from the same cell after it had been perfused for two minutes with KRB containing 1 μM nitrendipine (Alomone Cat# N-155). To isolate gating currents and *I*_tail_ produced by Cav1.2, we subtracted currents measured in the presence of nitrendipine from those measured in KRB alone. The on-gating charge (*Q*_on_) was then obtained from these records by integrating the gating current within approximately 2 ms of a depolarizing step to the reversal potential, and maximal *I*_tail_ amplitudes were measured upon repolarization to the holding potential.

Somatic whole-cell patch clamp recordings were acquired from WT and Kv2.1 KO mouse CHNs cultured on microscope cover glasses after 15-16 DIV. Pyramidal neurons were selected based upon their morphological characteristics (Benson et al., 1994). Patch pipettes were fashioned and filled with intracellular recording solution as described above. After establishing the whole-cell configuration in KRB, the bath solution was exchanged with an extracellular recording buffer containing (in mM): 135 NMDG, 30 TEA-Cl, 5 BaCl_2_, 8 glucose, 20 HEPES, adjusted to pH7.4 with HCl. Series resistance was 9.9±0.9 (WT) and 10.4±0.9 (Kv2.1 KO) MΩ (p=0.694, Student’s *t*-test) (before compensation); cell capacitance was 52.9±4.8 (WT) and 58.4±4.0 (Kv2.1 KO) pF (p=0.789, Student’s *t*-test). Prior to recording, cell capacitance was canceled, and series resistance was partially (60-70%) compensated. Recordings of LTCC ionic and gating currents were then performed as described for HEK293T cells. We used 10 µM nimodipine to isolate the contribution of LTCCs to the measured currents.

For simultaneous measurement of the *V*_m_ and Ca^2+^ sparks, rat CHNs transfected with GCaMP3-Kv2.1 were recorded using the whole-cell perforated patch clamp configuration. CHNs were patched in KRB using pipettes filled with a solution containing (in mM): 135 K-gluconate, 15 KCl, 5 NaCl, 1 MgCl_2_, 0.1 EGTA, 10 HEPES, pH adjusted to 7.22 using KOH, and amphotericin B (Millipore Cat# 171375) dissolved in DMSO and added at a final concentration of approximately 50 µg/mL. Upon obtaining a GΩ seal, the amplifier was switched to the current clamp mode to record spontaneous fluctuations in the *V*_m_. Measurement of the *V*_m_ (sampled at 25 kHz) and widefield image acquisition (acquired at 0.2 Hz) were triggered simultaneously using the same microscope described above.

### Experimental design and statistical analysis

For all data sets presented in this study for which statistical analysis were performed, measurements were imported into GraphPad Prism and Microsoft Excel for presentation and statistical analysis. Exact p-values are reported in each figure or figure legend. Proteomics on brain samples were collected from a from three independent sets of age-matched male wild-type and Kv2.1 KO adult mice. For experiments involving HEK293T cells and CHNs, at least two independent cultures were used for experimentation; the number of samples (*n*) indicates the number of cells analyzed and is noted in each figure legend.

## Acknowledgements

We thank Kimberly Nguyen and Grace Or Mizuno for assistance in preparation of rat CHNs. Much of the microscopy in this study was performed using the resources of the UC Davis MCB Imaging Facility. The proteomics experiments in this study were performed using the resources of the UC Davis Proteomics Core Facility. This project was funded by National Institutes of Health Grants U01NS0997 (JST), R21NS101648 (JST), R01HL144071 (LFS and JST), T32GM007377 (MK), and F32NS108519 (NCV).

---

**Figure S1.**
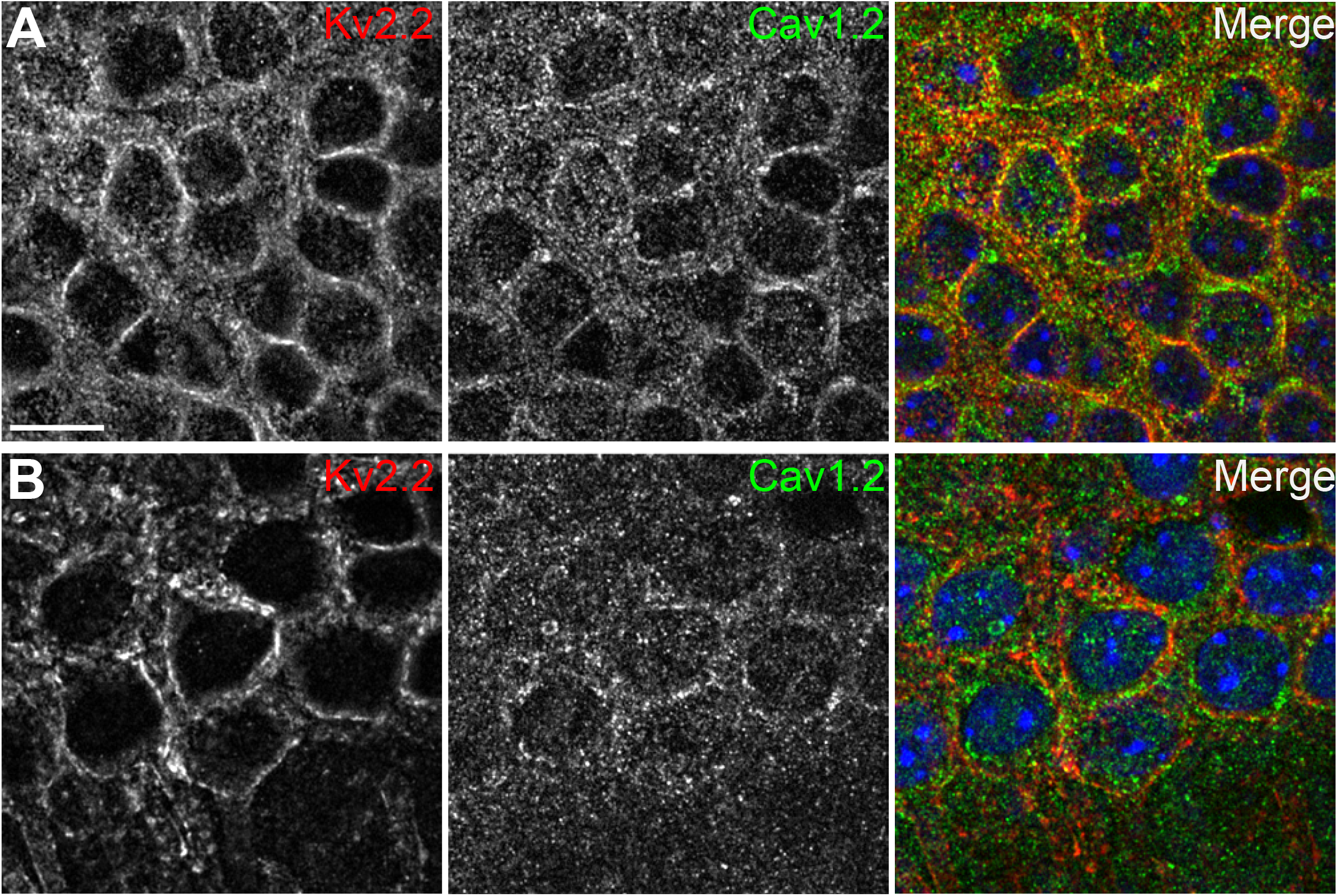
Cav1.2 spatially associates with Kv2.2 in brain neurons. (A) Confocal optical section obtained from the dentate gyrus of a rat brain section immunolabeled for Kv2.2 (red) and Cav1.2 (green). Nuclei are shown in blue (scale bar: 10 μm). (B) Confocal optical section obtained from the pyramidal cell layer of hippocampal area CA1 in a rat brain section immunolabeled for Kv2.2 (red) and Cav1.2 (green). Nuclei are shown in blue.

**Figure S2.**
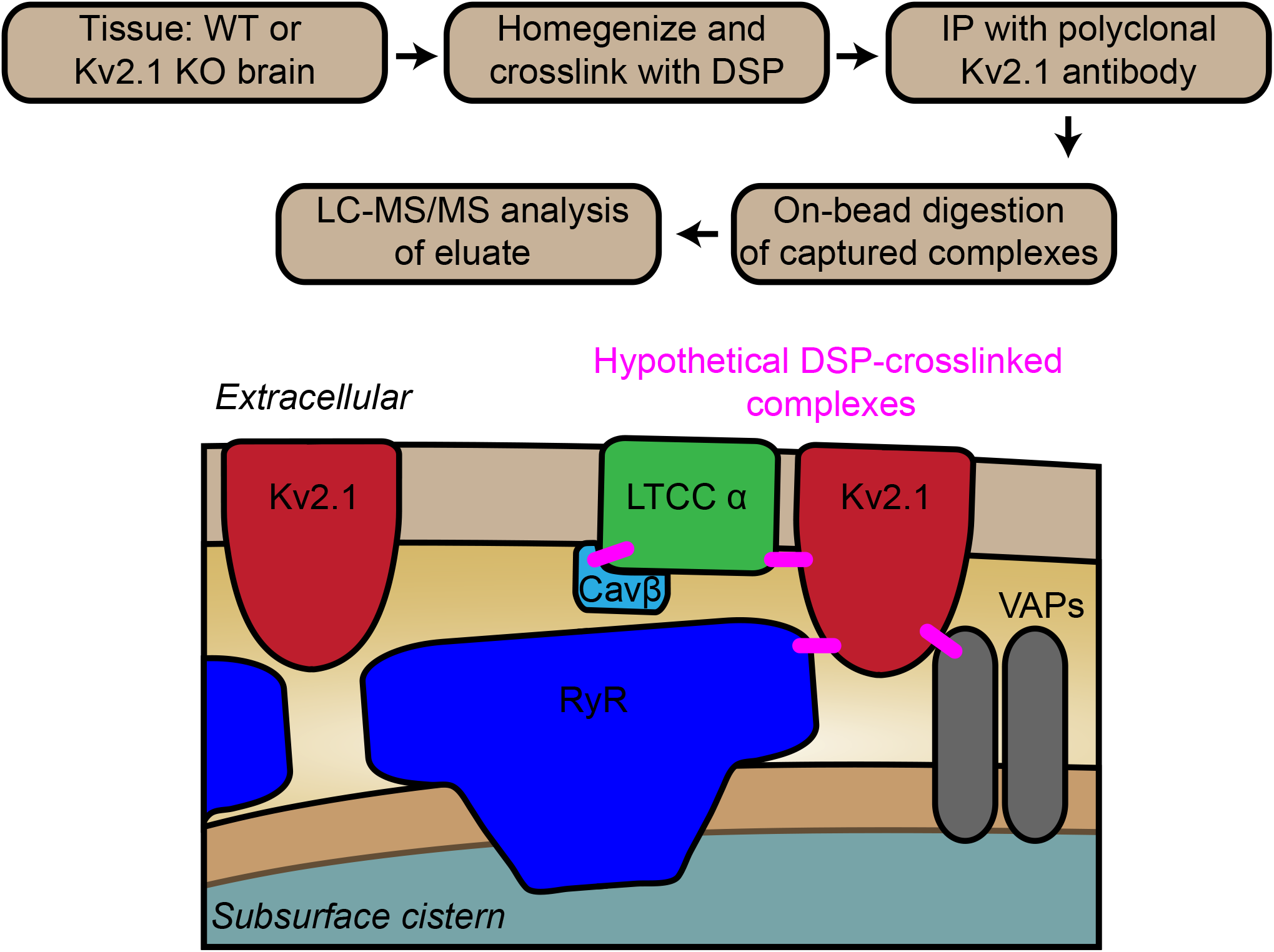
Identification of proteins in close spatial proximity to Kv2.1 using immunopurification and chemical crosslinking-based proteomics. Schematic detailing experimental workflow for immunopurification of proteins in close spatial proximity to Kv2.1 by chemical crosslinking of mouse brain homogenates, immunopurification of Kv2.1 and mass spectrometry (LC-MS/MS) based protein identification. Diagram below illustrates a model for the molecular architecture of Kv2.1-associated EPJs deduced from immunolabeling of neurons and crosslinking-based proteomics (note crosslinks between proteins shown in pink are hypothetical).

**Figure S3.**
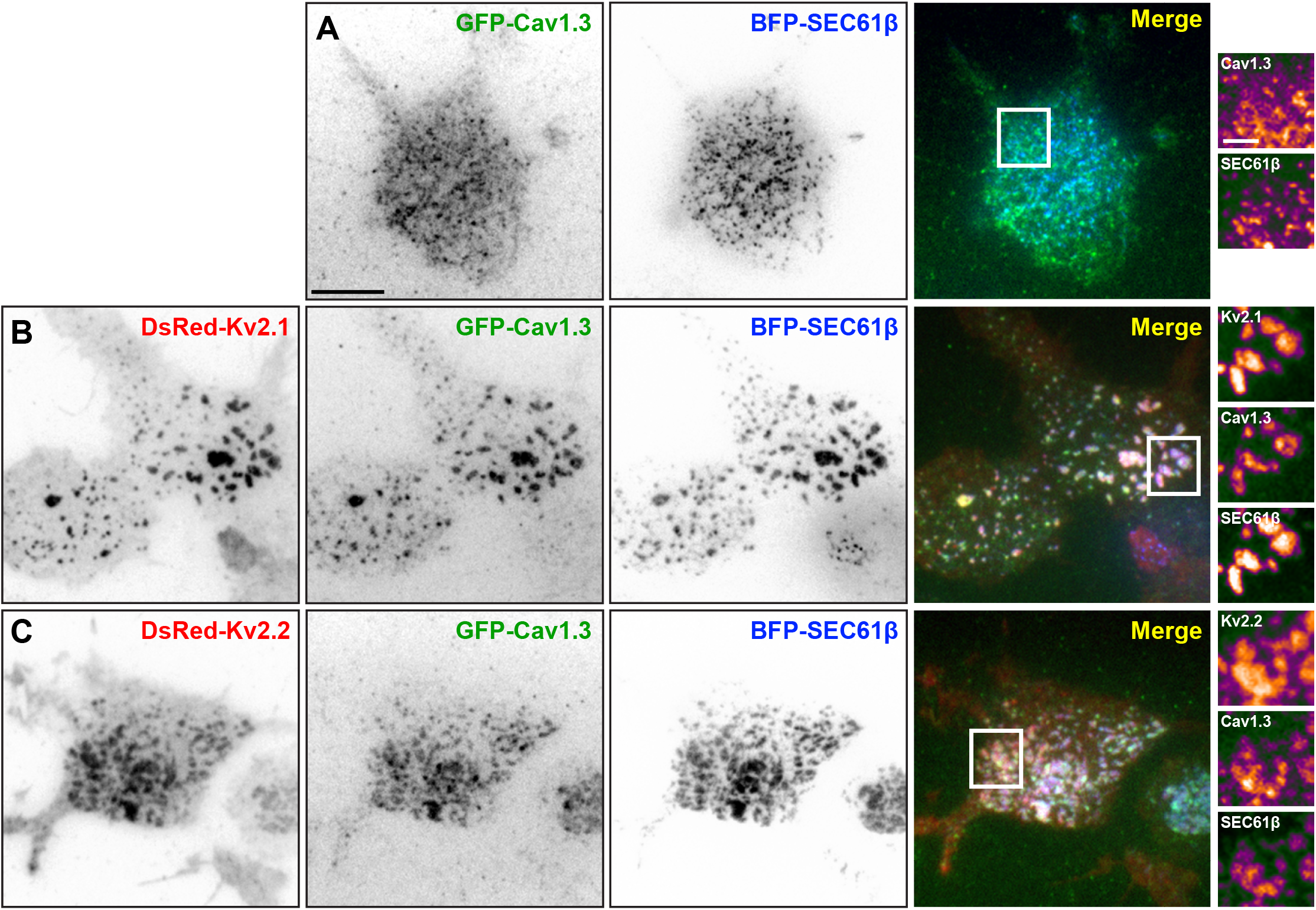
Cav1.3s is recruited to Kv2-induced EPJs. (A) TIRF images of a HEK293T cell cotransfected with the short isoform of Cav1.3 (GFP-Cav1.3 (green), BFP-SEC61β (blue) and auxiliary subunits Cavβ3 and Cavα2δ (not shown). Scalebar is 10 μm and holds for all large panels in figure. Pseudocolored intensity profiles of GFP-Cav1.3 and BFP-SEC61β, from the boxed area in the merged image, are shown to the right of merged image. (scale bar: 2.5 μm and holds for all pseudocolored intensity profiles in figure). (B) TIRF images of HEK293T cells cotransfected with DsRed-Kv2.1 (red), GFP-Cav1.3 (green), BFP-SEC61β (blue) and auxiliary subunits Cavβ3 and Cavα2δ (not shown). Pseudocolored intensity profiles of DsRed-Kv2.1, GFP-Cav1.3 and BFP-SEC61β, from the boxed area in the merged image, are shown to the right of merged image. (C) TIRF images of a HEK293T cell cotransfected with DsRed-Kv2.2 (red), GFP-Cav1.3 (green), BFP-SEC61b (blue) and auxiliary subunits Cavβ3 and Cavα2δ (not shown). Pseudocolored intensity profiles of DsRed-Kv2.2, GFP-Cav1.3 and BFP-SEC61β from the boxed area in the merged image, are shown to the right of merged image.

**Figure S4.**
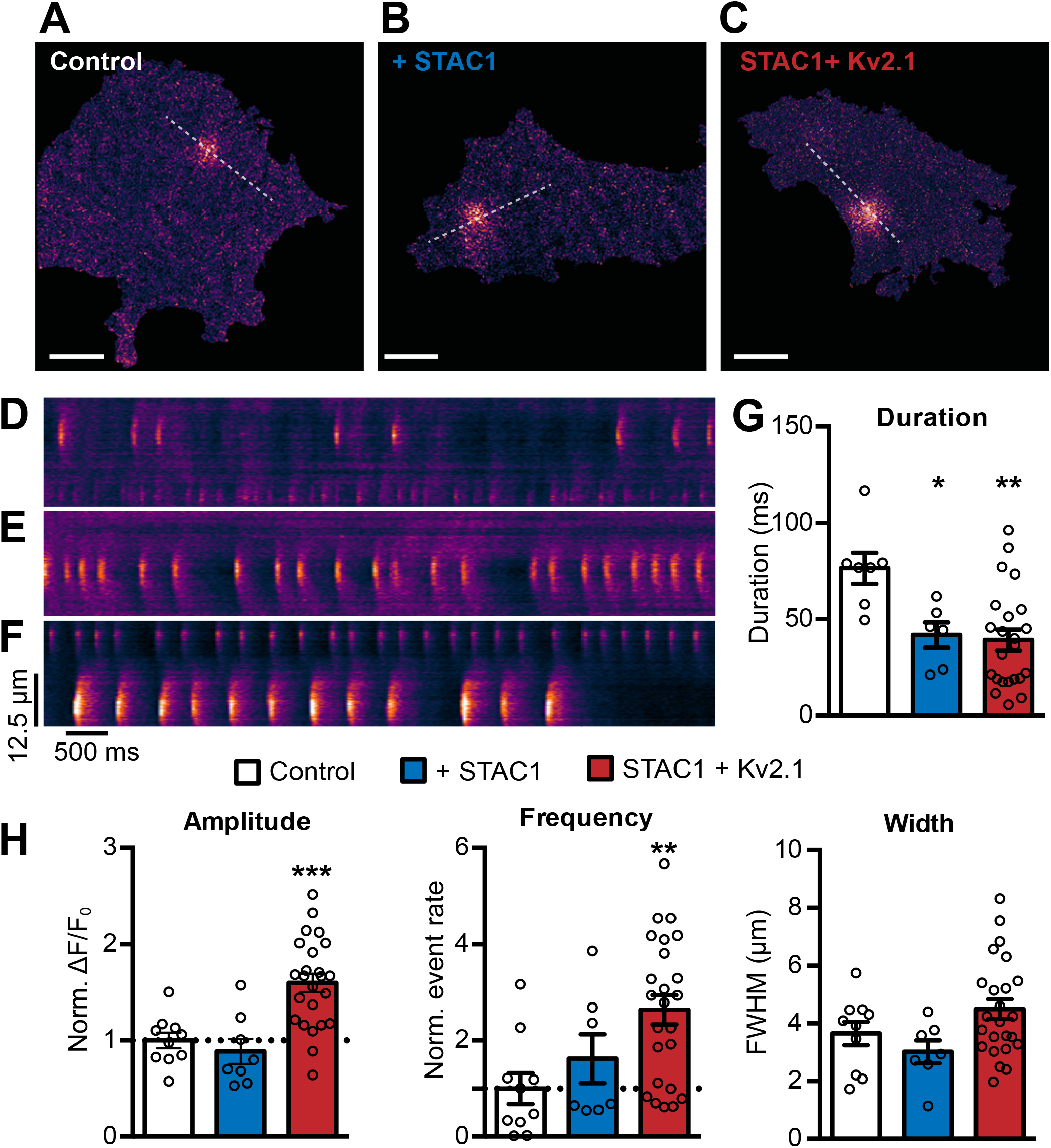
Kv2.1 increases the frequency of Cav1.3s and RyR-mediated sparks reconstituted in HEK293T cells. (A) TIRF image of HEK293T cell expressing the short isoform of Cav1.3 (Cav1.3s), RyR2, STAC1, and the LTCC auxiliary subunits β3 and α2δ1, and loaded with Cal-590 AM (scale bar: 10 µm and holds for panels A-C). (B) TIRF images of HEK293T cells additionally coexpressing STAC1. (C) TIRF images of HEK293T cells additionally coexpressing STAC1 and Kv2.1. (D-F) Kymographs showing the localized Ca^2+^ release events detected in the ROI on the cell in panels A-C, respectively. (G-H) Data from cells expressing Cav1.3, RyR2 and auxiliary subunits without (white bars) or with coexpression of Stac1 (blue bars) or Stac1 + Kv2.1 (red bars). (G) Expression of STAC1 reduces the duration of Cav1.3s- and RyR2-mediated CICR events reconstituted in HEK293T cells. (*p=0.0339; **p=0.0026; ANOVA followed by Dunnett’s test). (H) Summary data of the amplitude, frequency, and spatial spread (width) of all sparks recorded. Each point corresponds to a single cell (**p=0.0081; ***p=0.0001; ANOVA followed by Dunnett’s test).

**Figure S5.**
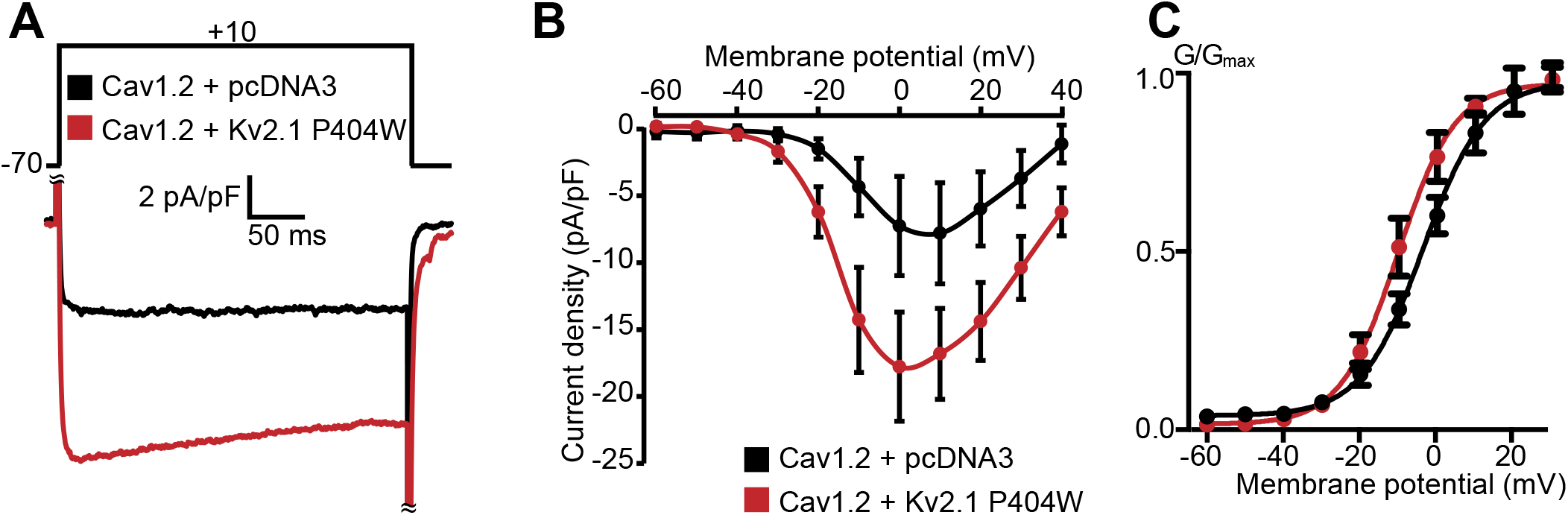
Cav1.2 channel activity is increased in cells coexpressing Stac1 upon coexpression with Kv2.1_P404W_. A-C: Data recorded from HEK293T cells transfected with Cav1.2-GFP and auxiliary subunits Cavβ3, Cavα_2_δ_1_, and STAC1, without (+pCDNA3, in black) or with Kv2.1_P404W_ (in red). (A) Representative Ca^2+^ current traces at +10 mV. (B) Normalized *I-V* relationship of whole-cell *I*_Ca_ recorded from *n*=8 (Cav1.2 + pcDNA3) and *n*=9 (Cav1.2 + Kv2.1_P404W_) cells. (C) Voltage-dependence of whole cell Cav1.2 conductance *G*/*G*_max_. For the conductance-voltage relationships, the half-maximal activation voltage *V*_1/2_=1.6±2.0 [pcDNA3] vs. -9.5±2.9 [+Kv2.1_P404W_] mV, p=0.0166; slope factor k=8.8±1.2 [pcDNA3] vs. 6.1±0.6 [+Kv2.1_P404W_], p=0.0490; Student’s *t*-test).

**Movie S1. Spontaneous somatic Ca^2+^ signals detected at GCaMP3-Kv2.1 clusters in cultured rat CHNs**. Stack of widefield images of a rat CHN transfected with GCaMP3-Kv2.1 and imaged at 10 Hz.

**Movie S2. Spontaneous somatic Ca^2+^ signals detected by TIRF microscopy in cultured rat CHNs loaded with Cal-590 AM**. Stack of TIRF images of rat CHNs loaded with Cal-590 AM and imaged at 30 Hz. Regular wave-like signals are a TIRF imaging artifact. Images have been normalized to the first image without detectable Ca^2+^ signals (i.e., F/F_min_).

**Movie S3. Caffeine increases the frequency of somatic Ca^2+^ sparks in cultured CHNs**. Images of a rat CHN transfected with GCaMP3-Kv2.1 acquired at 5 Hz. Neuron is treated with 5 mM caffeine at t=84 s; the increased Ca^2+^ spark frequency is apparent from t=87s-101s. Images have been normalized to the first image without detectable Ca^2+^ signals (i.e., F/F_min_).

**Movie S4. Bay K8644 increases the frequency of somatic Ca^2+^ sparks in cultured CHNs**. Rat CHN transfected with GCaMP3-Kv2.1 and imaged in the presence of 500 nM Bay K8644. Images have been normalized to the first image without detectable Ca^2+^ signals (i.e., F/F_min_).

**Movie S5. Tetracaine blocks Ca^2+^ sparks reconstituted in HEK293T cells**. Stack of TIRF images of a single HEK293T cell transfected with RyR2, Cav1.2, and auxiliary subunits and loaded with Cal-590 AM. 100 µM tetracaine was added at t=7000 ms. Regular wave-like signals are a TIRF imaging artifact. Images have been normalized to the first image without detectable Ca^2+^ signals (*i.e.*, F/F_min_).

